# Mouse Model Systems of Autism Spectrum Disorder: Replicability and Informatics Signature

**DOI:** 10.1101/561233

**Authors:** Patricia Kabitzke, Diana Morales, Dansha He, Kimberly Cox, Jane Sutphen, Lucinda Thiede, Emily Sabath, Taleen Hanania, Barbara Biemans, Daniela Brunner

**Author notes:** These authors contributed equally to this work (Patricia Kabitzke, Daniela Brunner).

## Abstract

3.

**Background:** Phenotyping mouse model systems of human disease has proven to be a difficult task, with frequent poor inter- and intra-laboratory replicability and translatability, particularly in behavioral domains such as social and verbal function. However, establishing robust animal model systems with strong construct validity is of fundamental importance as they are central tools for understanding disease pathophysiology and developing therapeutics. To complete our studies of mouse model systems relevant to autism spectrum disorder (ASD), we present a replication of the main findings from our two published studies comprising five genetic mouse model systems of ASD.

**Methods:** To assess the robustness of our previous results, we chose the two model systems that showed the greatest phenotypic differences, the *Shank3/F* and *Cntnap2*, and repeated assessments of general health, activity, and social behavior. We additionally explored all five model systems in the same framework, comparing all results obtained in this three-yearlong effort using informatics techniques to look for commonalities and differences.

**Results:** Results in the current study were very similar to our previously published results. The informatics signatures of the two model systems chosen for the replication showed that they were most distinguished by activity levels. Although the two model systems were opposite in this regard, those aspects of their social behavior not confounded by activity (vocalizations) were similar.

**Conclusions:** Our results showed high intra-laboratory replicability of results, even for those with effect sizes that were not particularly large, suggesting that discrepancies in the literature may be dependent on subtle differences in testing conditions, housing enrichment, or background strains and not so much on the variability of the behavioral phenotypes. The overall informatics analysis suggests two main classes of model systems that in some aspects lie on opposite ends of the behavioral spectrum, supporting the view that autism is not a unitary concept.

## 5. Main Body

### 5.1. Introduction

Autism spectrum disorder (ASD) has been linked to gene copy number and single nucleotide variation, findings that lead to a number of mouse model systems with etiological validity (Moy and Nadler 2008, Ey, Leblond et al. 2011, Jiang and Ehlers 2013, Kazdoba, Leach et al. 2016). We previously completed and published two studies from a considerable effort using standard and informatics methods to phenotype five mouse model systems of ASD. The first study on the 16p11.2 and *Cntnap2* deletion mouse model systems (Brunner, Kabitzke et al. 2015), and the second complementary study on two distinct *Shank3* knockout (KO) mouse model systems of Phelan-McDermid Disorder, Feng’s *Shank3*^*tm2Gfng*^ (hereafter *Shank3/F*) and Jiang’s *Shank3^tm1Yhj^* (hereafter *Shank3/J*), and a model of Timothy syndrome, the *Cacna1c* heterozygous (HET) mouse model system (Kabitzke, Brunner et al. 2018). In the current study, we focus on two separate questions. Can we replicate the main results of our published studies? Do the 5 model systems lie on a behavioral continuum, or do they present idiosyncratic signatures?

#### Replication and robustness

Numerous papers, including our own (Brunner, Balci et al. 2012, Golani, Wexler et al. 2014, Munafo, Noble et al. 2014, Begley and Ioannidis 2015, Brunner, Kabitzke et al. 2015, Peng 2015, Jarvis and Williams 2016, Kabitzke, Brunner et al. 2018), have pointed at the difficulty of replicating results from preclinical and clinical studies, and offer various explanations including the likelihood of false positives due to small sample sizes. In our previous publications, we chose to present all the results of our broad battery of behavioral endpoints and their independent analysis. Rather than using statistical corrections to reduce the experiment-wise Type I error, we chose to do a real replication of the main findings. This route ensures fair assessment of effect sizes that may not be large yet may be of high scientific interest, which would be obliterated by simple statistical corrections.

It should be noted too that *p*-value corrections should only be applied to results that are considered to be of the same “family”, that is, tests that assess the likelihood of falsehood of the same hypothesis (Cabin and Mitchell 2000). For example, within a phenotyping project, one could use two different measures of activity to assess if the model system has an abnormal motor pattern as compared to its control. These two measures are testing the same hypothesis, namely, “does this mouse model exhibit abnormal motor activity”. Other aspects of the phenotype (e.g. social, cognitive, etc.) should be considered under other null hypotheses. It could be parsimonious, therefore, to assume that these other aspects of the phenotype have different underlying biological underpinnings better tested under independent hypotheses, which would not belong to the same “hypothesis family”.

It is also important to differentiate between an exploratory analysis, such as a broad phenotyping effort, and a focused experimental exercise, such as a drug test for which a primary endpoint measure should be defined *a priori* (Kimmelman, Mogil et al. 2014). Exploratory analyses should be taken as tentative poking of the phenotypical landscape to find where the signal resides and should be followed by replication. Thus, replication rather than statistical corrections should be the rule, contrary to the common practice to take single studies as proof of a phenomenon (Button, Ioannidis et al. 2013). Taking this to heart, we bred independent groups of mice, from the *Shank3/F* and *Cntnap2* model systems and chose to replicate phenotypic differences found in our published studies (Brunner, Kabitzke et al. 2015, Kabitzke, Brunner et al. 2018). The SmartCube^®^ platform provided a comprehensive analysis. This test together with the reciprocal social interaction test and the urine-exposure open field test assessed behavior in a social setting. We added the standard open field test, not done before, to provide an assessment of activity levels in a non-social setting. Body weight measures helped to probe general health and our ability to replicate our own results.

#### ASD signatures: Common Elements or Idiosyncratic Features?

A comprehensive analysis of behavior provides a panorama of complex function reflecting the downstream effects of the genetic insult or manipulation. Phenotypic analyses of model systems of ASD typically probe mouse functional domains putatively reflecting the core ASD triad, namely, social function, communication, and repetitive behaviors (Moy, Nadler et al. 2006, Crawley 2007, Happé and Ronald 2008, Silverman, Yang et al. 2010). In addition to those standard tests, we previously used additional proprietary platforms, SmartCube^®^ and NeuroCube^®^, to discover possible unexpected phenotypes. SmartCube^®^ captures ½ million points related to posture, position, trajectory, activity, behavioral sequences, response to stimulation, and individual behaviors. This dataset is reduced to a several hundred informative features, constituting a rich content dataset amenable to machine learning mining techniques (Brunner, Nestler et al. 2002, Tecott and Nestler 2004, Alexandrov, Brunner et al. 2015). We took advantage of these features to run a comprehensive analysis of the results from the original SmartCube^®^ studies covering the five model systems and added the new results from the tests presented in this paper. We asked the following questions: Do the model systems lie in a complex plane or could their similarities and differences be well explained using a few dimensions? Can we classify one mouse as being a mutant or a wild type by training the classifier on a different model system? Indeed, which model systems are confused with each other and which are distinct? Is classification accuracy particularly high when we attempt to classify the replicates, having trained on the original dataset? These informatics exercises attempt to find core characteristics of the mutant mice that may be relevant to ASD-defining features in an agnostic and unbiased way.

### 5.2. Methods

#### Ethics Statement

PsychoGenics is an AAALAC accredited facility (Unit Number – 001213) and work is performed under PHS OLAW Assurance #A4471-01. This study was carried out in strict accordance with the recommendations in the Guide for the Care and Use of Laboratory Animals of the National Institutes of Health. The protocol was approved by the Committee on the Ethics of Animal Experiments of PsychoGenics. All efforts were made to minimize suffering and maximize animal welfare.

#### Subjects

##### *Shank3/F* Line

###### Breeders

A cohort of 60 Het female and 30 male mice (JAX 3258240; stock 017688, B6.129- *Shank3*<tm2Gfng>/J) was provided by The Jackson Laboratory at 11-13 weeks of age. Line development prior to arrival at JAX: Exons 13-16 of SH3/ankyrin domain 3 were replaced with a neomycin resistance (neo) cassette. The construct was electroporated into (129X1/SvJ x 129S1/Sv)F1-Kitl^+^-derived R1 embryonic stem (ES) cells. Correctly targeted ES cells were injected into C57BL/6 blastocysts and the resulting chimeric males were bred to C57BL/6J females. The offspring were then intercrossed for five generations and maintained on a mixed C57BL/6J x 129 background prior to sending to The Jackson Laboratory. Line maintenance at JAX: Upon arrival, mice were additionally backcrossed to C57BL/6J inbred mice (Stock No. 000664) using a marker-assisted, speed congenic approach to establish this congenic line. Genome Scan results indicated that *Shank3* Feng breeders were fully congenic for C57B6J.

##### *Cntnap2* Line

###### Breeders

A cohort of 60 female and 30 male *Cntnap2* -/- (B6.129(Cg)-Cntnap2^tm1Pele^/J) mice (catalog #017482), backcrossed to C57BL/6J for more than 10 generations, was provided by The Jackson Laboratory at 8-13 weeks of age.

##### Breeding Scheme

Mice were set in trios (2 females : 1 male) and left together for three days. Breeders were 14-16 weeks of age when bred for the *Shank3/F* line and 11-16 weeks of age when bred for the *Cntnap2* line. See S1 Methods for genotyping results. Sex and genotype ratios were unbiased in all animal model systems. Animals were weaned at 4 weeks of age. Breeding success and pup survival were similar among the four groups. All testing was done in male mice. Mutants and their wild type controls were littermates. Age-matched male (∼P45) unfamiliar littermates were used as stimulus mice for the reciprocal social interaction test. Stimulus mice were used for the Grooming test to maintain the same sequence of testing used in other studies, where tests were performed longitudinally in the same animals.

#### General Procedures

Mice were housed in OptiMice cages (Animal Care Systems, Inc.) on a 12/12hr light/dark cycle where 20–23°C room temperature and a relative humidity of 50% were maintained. Chow and water were provided ad libitum for the duration of the study and mice were checked twice daily for general health, food, and water. Husbandry included enrichment, namely, shredded paper (Enviro-Dri; W.F. Fisher & Son Inc., NJ; Product 08ENV3000) and a nylabone (Bio-Serv, NJ; Product K3200). Breeders also received an amber-colored polycarbonate igloo for extra enrichment (Bio-Serv, NJ; Product K3328). On P0, pups were tattooed using non-toxic ink applied under the skin of their toe and a tail snip sample was taken for genotyping (see S1 Methods). Once the genotype results were available (around P2), the litter size was culled down to N = 8 pups, removing mainly females via decapitation. Thus, litter size range was (3–8) after culling. Animals were weaned in 2:2 mixed-genotype, same-sex groups of four with shredded paper, one nylabone, and one polycarbonate amber-colored tunnel (3 7/8” long x 2” inside diameter) (Bio-Serv, NJ; Product K3323) per cage. Testing occurred between 10:00 and 17:30 in separate experimental rooms. Tests were conducted blind to genotype and the sequence of testing is indicated in Table 1. Euthanasia is required at first signs of illness, severe dehydration and/or emaciation defined as a loss of greater than 20% body weight with failure to regain weight while on a free feeding regimen, lack of righting reflex, catalepsy, morbidity, increased repetitive convulsions, respiratory distress, or hemorrhage. Although no mice were euthanized for any of these reasons, mice were sacrificed at the end of the study using methods consistent with recommendations of the 2013 American Veterinary Medical Association (AVMA) Guidelines on Euthanasia. Carbon dioxide gas was used and euthanasia was verified by observation of breathing and color of the animal, and by palpation of the heart in addition to the loss of reflexes. Required further verification of death was accomplished via cervical dislocation.

**Table 1.**
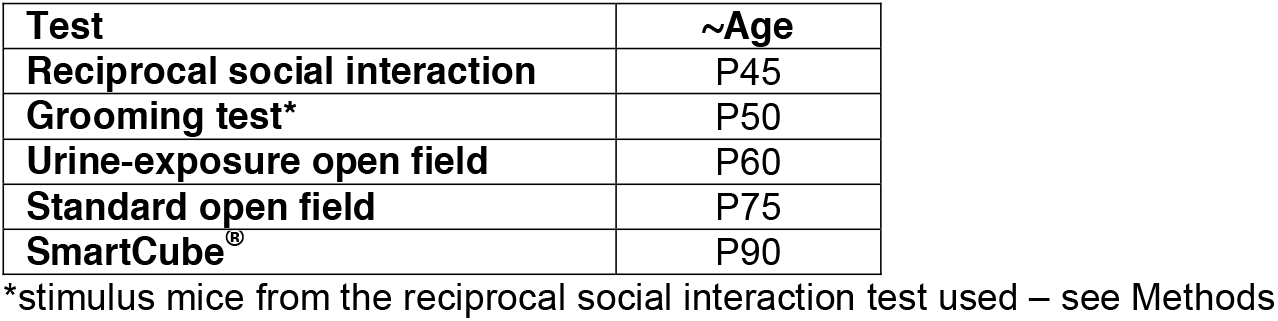
Time-Course of Tests.

##### Reciprocal Social Interaction Test

Subject animals were isolated for 2 days before testing. The day before testing, subject animals were individually habituated to the testing apparatus for 10 min and same-genotype stimulus animals (that were not of the same litter, were unfamiliar, and age-matched) were separately habituated to the apparatus in pairs. The day of testing, subject animals were placed in the testing apparatus for 5 min before a genotype-, age-, and weight-matched stimulus animal was placed into the chamber. Behavior and USVs were recorded for the pair for a total of 10 minutes. Ethovision XT (Noldus Information Technology, Wageningen, Netherlands) was used to measure distance, proximity, and interaction between animals. We defined close proximity as the center of the bodies being between 1 and 5 cm apart and interactions when the distance from the nose to the other mouse body (nose, center, and tail) was less than 1.5 cm. Interactions of the mice that were active (one mouse sniffing any part of the other mouse body), passive (recipient of the other subject’s investigation) and reciprocal (both mice actively sniffing each other) were scored manually for the first 5 minutes of the test period.

##### Grooming Test

The grooming test is used to measure anxiety and repetitive behavior during the first two hours of the dark cycle. Mice were individually housed in standard mouse cages for one week before the test. One hour prior to the test, at 5:30 pm, they were placed in the testing room to habituate. At 6:50 pm, mice were transferred to a standard cage with clean bedding, red lights were turned on and white room lights were turned off and the testing cage was recorded for 2 hours. After 2 hours of videotaping, animals were group housed in their original configurations and returned to the colony room.

##### Urine-Exposure Open Field Test

Procedures were based on those described in the literature (Wohr, Roullet et al. 2011). One week before the test, males were exposed to same-strain females for 5 minutes in a novel cage with fresh bedding. The day before the test a handful of soiled male bedding was placed in the female cages to induce estrus. Estrus was determined by visual inspection of the vaginal area. The open field was conducted in a dimly lit room. The adult male mice were placed in a clean open field, lined with paper (Strathmore Drawing Paper Premium, recycled, microperforated, 400 series; Strathmore Artist Papers, Neenah, WI, USA) and containing some of its own home cage bedding in a corner of the arena. Open field activity was recorded for 60 min. At the end of the habituation period, the mouse was placed back in a clean polycarbonate cage with fresh bedding. The home cage bedding and any feces deposited by the mouse were removed from the open field. Urinary scent marks deposited on the paper during habituation were visualized under ultraviolet (UV) light and outlined with a pencil for subsequent quantification. Fifteen microliters of fresh female urine, pooled from 4–6 estrous females, was then pipetted onto the center of the Strathmore paper, and the mouse was placed back into the open field for 5 min. Open field activity and ultrasonic vocalizations were recorded. The marked sheets of Strathmore paper were treated with Ninhydrin spray (LC-NIN-16; TritechForensics, Inc., Southport, NC, USA) then left to dry for ∼12 hours, which allowed the visualization of the urine traces as purple spots. Once dry, images were scanned and opened in ImageJ (U. S. National Institutes of Health, Bethesda, Maryland, USA). Freehand selections of the circled areas (pre-exposure marking) were removed and copied into a new JPEG image. The pre-exposure and post-exposure images were processed in 8-bit, with background subtracted, and converted to binary. Particles were analyzed at 1000-Infinity (pixels) and 0.00–1.00 (circularity), counted, and their area measured.

##### Standard Open Field Test

The open field test is used to assess motor activity. The open field chambers are Plexiglas square chambers (27.3 × 27.3 × 20.3 cm; Med Associates Inc., St Albans, VT) surrounded by infrared photobeam sources (16 × 16 × 16). The enclosure is configured to split the open field into a center and periphery zone and the photocell beams were set to measure activity in the center and in the periphery of the open field chambers. Horizontal activity (distance traveled) and vertical activity (rearing) are measured from consecutive beam breaks. Animals are placed in the open field chambers for a 60 min session and returned back to the home cages after test completion.

##### SmartCube^®^ Test

To enable phenotyping of mouse models of human disorders and testing of compounds for behavioral effects relevant to psychiatric disease, PsychoGenics developed SmartCube, an automated system in which mouse behavior is captured by digital video using novel, proprietary hardware that presents multiple challenges in a test session and is analyzed with computer algorithms. Digital videos of the subjects are processed with computer vision algorithms to extract more than 1400 dependent measures including frequency and duration of behavioral states such as grooming, rearing, etc., and many other features obtained during the test session. Using machine learning techniques chosen to best separate pharmacological effects of reference compounds, the behavioral signatures of the mutant mice are then assessed quantitatively (Brunner, Nestler et al. 2002, Houghten, Pinilla et al. 2008, Roberds, Filippov et al. 2011, Alexandrov, Brunner et al. 2015).

Mice were taken in their home cage to the SmartCube suite of experimental rooms where they remained until they were placed in the apparatus. A standard SmartCube protocol runs for a single session (∼45 min). After the session mice were group-housed again and brought back to the colony room. Any abnormal behavior was noted.

#### Data Handling

For all tests, unless noted otherwise, statistical analyses consisted of one- or two-way ANOVAs (StatView for Windows Version 5.0.1, SAS Institute Inc., Cary, NC) with Genotype as a between-subjects factor and, when appropriate, Session, or Stimulus type as a within-subject factor. Significant interactions between within- and between-subject factors were followed by simple main effects (SPSS, IBM). The level of significance was set at p < .05. No outliers were removed. For repeated measures ANOVAs, the data of a subject was removed when data was missing for such subject at a time point. Categorical data were analyzed with Mann Whitney and frequency data were analyzed with Chi-Square or Fisher Exact as noted.

##### Bioinformatics for SmartCube®

The most dominant of the features collected that define the phenotype (symptom descriptors) were identified and ranked using complex proprietary bioinformatics algorithms and an overall discrimination index was calculated for all features combined or for different subsets of features. Graphical representations of the datasets corresponding to the groups compared were derived and a p-value was calculated to assess the statistical significance of the discrimination ratios. Top representative features were graphically presented to aid interpretation of differences (see S1 Methods) (Alexandrov, Brunner et al. 2015).

###### Feature analysis: De-correlation and ranking

The outcome of a SmartCube run is a set of ∼1400 features (behavioral parameters) that can be used for various analyses. Many of these features are correlated (e.g. rearing counts and supported rearing counts). Therefore, we form statistically independent combinations of the original features (further referred to as de-correlated features) that discriminate between the two groups more effectively. Each de-correlated feature extracts information from the whole cluster of the original features, so the new feature space has lower dimensionality. Next, we apply a proprietary feature ranking algorithm to score each feature discrimination power (ability to separate the two groups, e.g. control and disease). Ranking is an important part of our analyses because it weighs each feature change by its relevance: if there is a significant change in some irrelevant feature measured for a particular phenotype, the low rank of this feature will automatically reduce the effect of such change in our analyses, so we do not have to resort to the conventional “feature selection” approach and discard information buried in the less informative features. The ranking algorithm can be applied to either original or the new features to gain insight about the key control-disease differences.

###### Feature analysis: Quantitative assessment of disease phenotype

In the new decorrelated feature space, the overlap between the “clouds” (Gaussian distributions approximating the groups of mice in the ranked de-correlated features space) serves as a quantitative measure of separability (“distinguishability”) between the two groups. For visualization purposes, we plot each cloud with its semi-axes equal to the one standard deviation along the corresponding dimensions. Note, however, that while the overlap between any two Gaussian distributions is always non-zero, it may not necessarily be seen at the “1-sigma level”. As in over-determined systems, the discrimination index sometimes can be artificially high, we calculate its statistical significance by estimating the probability that the result is due to chance.

###### Top features identification

Working back from the discrimination analysis we can identify the features that contribute the most to the separation between the two groups. Although statistical significance for differences between groups for the individual top features can be calculated, the alpha value for such statistical exercise cannot be set to the standard p = .05, as dozens of features are measured and combed for differences. Instead of over-interpreting such top features we present them in order to understand the mutant signatures but refrain from performing misleading standard statistical tests.

### 5.3. Results

#### Intralaboratory Replication

##### SmartCube®

SmartCube^®^ is a high-throughput automated behavioral platform that provides a comprehensive phenotypical assessment of mouse disease model systems and drug effects. Through computer vision, mechanical actuators, and machine learning techniques it identifies individual behaviors, postures, positions, trajectories, sequences of behaviors and responses to stimulation (Brunner, Nestler et al. 2002, Houghten, Pinilla et al. 2008, Roberds, Filippov et al. 2011). Machine learning analysis is also used to identify the features within the many hundreds measured (see S1 Methods) that contribute the most to the separation of classes under study. We plot data in the 2D space that best separates mutant from wild type groups, using the overlap between groups as a discrimination index. Each mutant group was compared against its own wild type (WT) group, with independent classifiers being trained for each mutant model against its corresponding WT group.

In the replication study, the discrimination index between *Shank3/F* WT and KO mice reached 95% (Fig 1A). The probability that this discrimination value could be obtained by chance is negligible (Fig 1B). Top features contributing to this difference were reduced vertical movements, decreased startle, increased freezing, decreased sniffing and digging (Fig 3 and Fig 4; S1 Table). The discrimination between the *Cntnap2* -/- and WT mice reached 97% also with a very small *p*-value (Fig 2A and 2B). The top features for this model were somehow opposite: hyperactivity, higher speed, increased movements, more missteps, and increased startle (Fig 3 and Fig 4; S1 Table). Figure 1 and 2 show the previous original study results with 74% and 88% discriminations for the *Shank3/F* and *Cntnap2* model systems, respectively.

**Fig 1.**
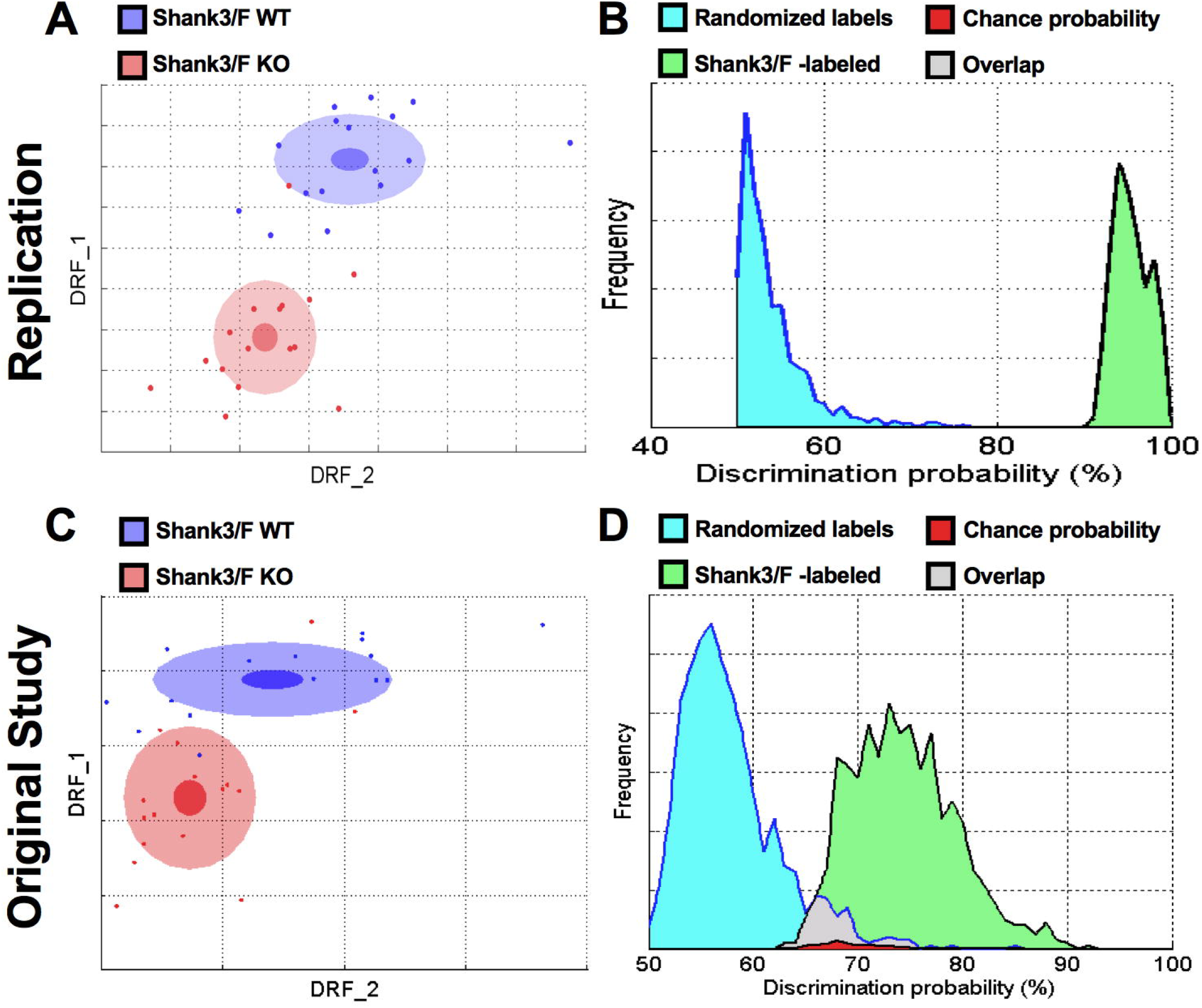
*Shank3/F* KO and WT littermates were very different in the SmartCube^®^ test across both studies. A & C: 2D representation of the multidimensional space in which the two groups are best separated. The DRF1 and 2 axes represent statistically independent combinations of the raw features that best discriminate between the two groups whereas the overlap between the groups is a measure of discriminability (S1 Methods). The dots represent individual mice (blue: WT; red: KO mouse). The center, small and large ellipses are the mean, standard error and standard deviation for each group, respectively. B & D: Discrimination indexes are repeatedly calculated, and their distribution plotted, using either correct labels (green distribution) or randomized labels (blue distribution) such that the overlap between the two distributions (in red) represents the probability of obtaining the observed discrimination by chance. A-B: In the replication study, the *Shank3/F* model separated well from the WT group with 95% discrimination and *p*< 0.0001. C-D: In the original study, the *Shank3/F* groups separated with 74% discrimination with *p*= 0.02. N=16 mice per genotype/line (replication); n=15-16 mice per genotype/line (original study).

**Fig 2.**
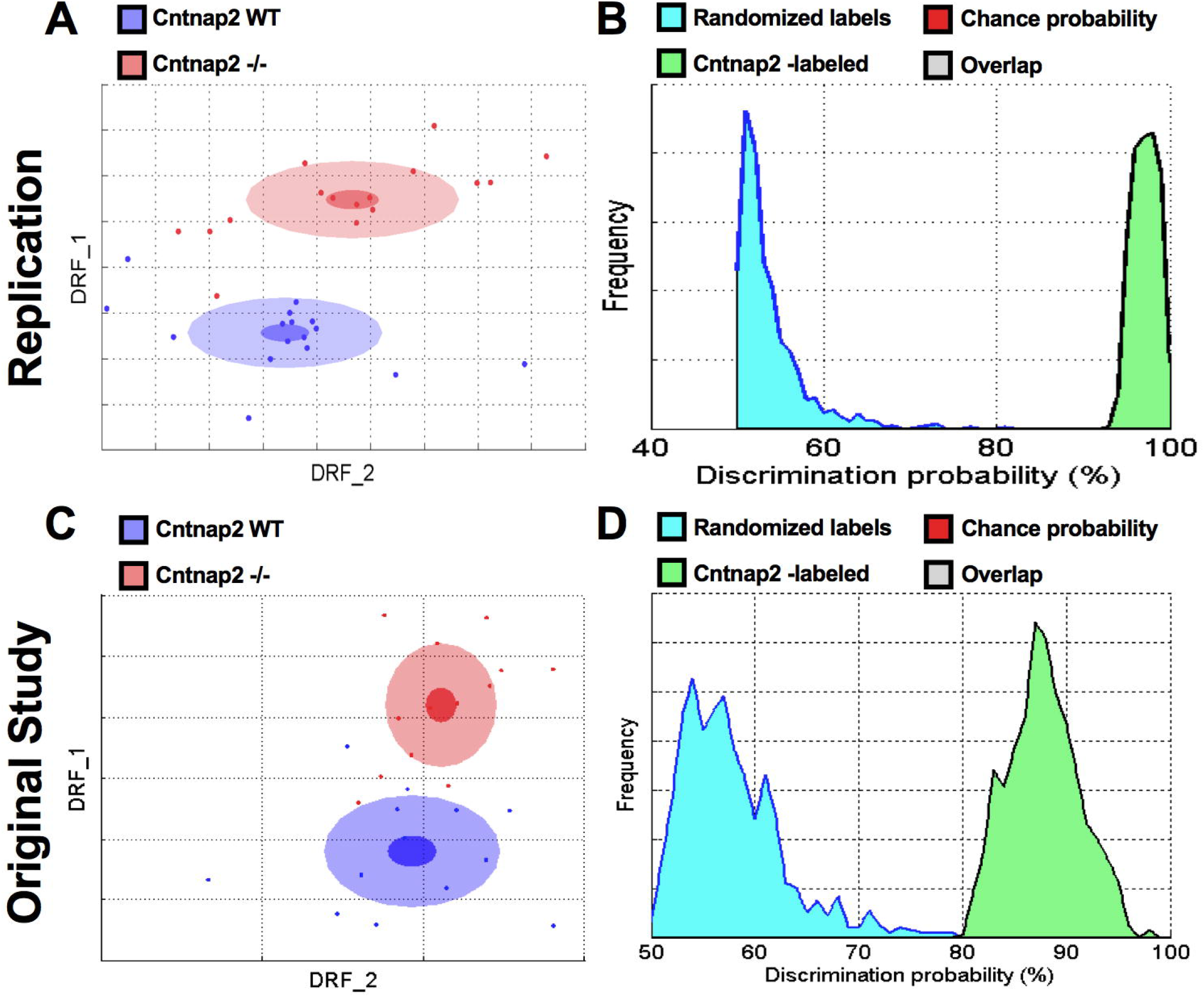
*Cntnap2 -/-* and WT littermates were very different in the SmartCube^®^ test across both studies. A-B: In the replication study, the *Cntnap2* model separated well from the WT group with a 97% discrimination and *p*< 0.0001. C-D: In the original study, the *Cntnap2* groups separated with 88% discrimination with *p*< 0.0002. N=16 mice per genotype/line (replication); n=13 mice per genotype/line (original study).

**Fig 3.**
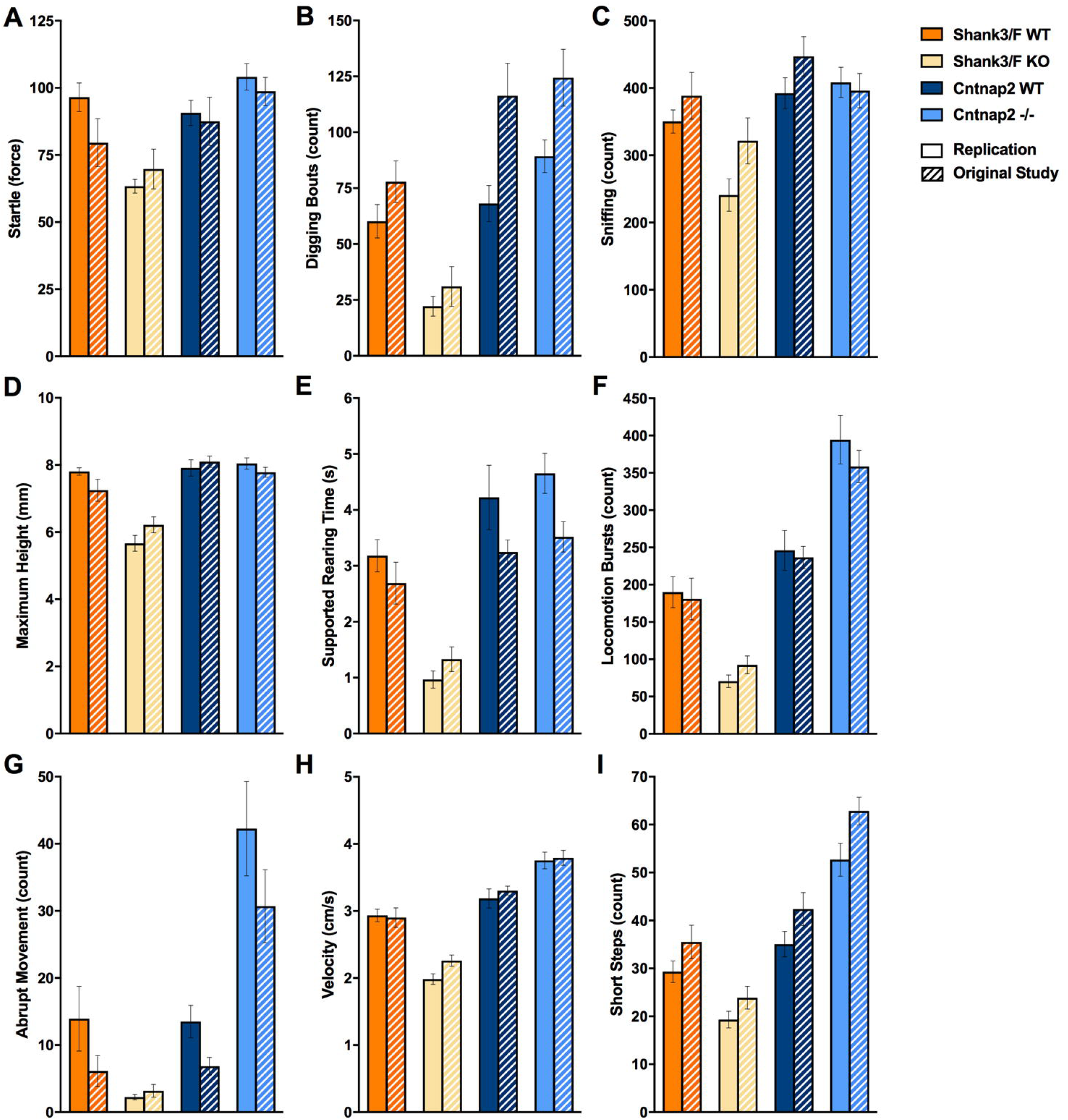
Top features in SmartCube^®^ showed changes in opposite directions for the models across both studies. The top features that separate the mutant and control groups are plotted for both models and for the replication (filled bars) and original (hashed bars) study. Remarkably, even with more than a year between the original and replication study, most of the results were extremely similar. Across models, many features showed decreases in the *Shank3/F* KO mice compared to corresponding WT mice and increases or no effect in the *Cntnap2* -/-mice compared to WT littermates. *Shank3/F* KO mice showed (or tended to show) decreased startle (A), digging (B), sniffing (C), average height (D), rearing (E), locomotion (F), abrupt movements (G), velocity (H), and short steps (I) compared to WT littermates, whereas the *Cntnap2* -/- model showed no change or increases in measures compared to WT littermates.

**Fig 4.**
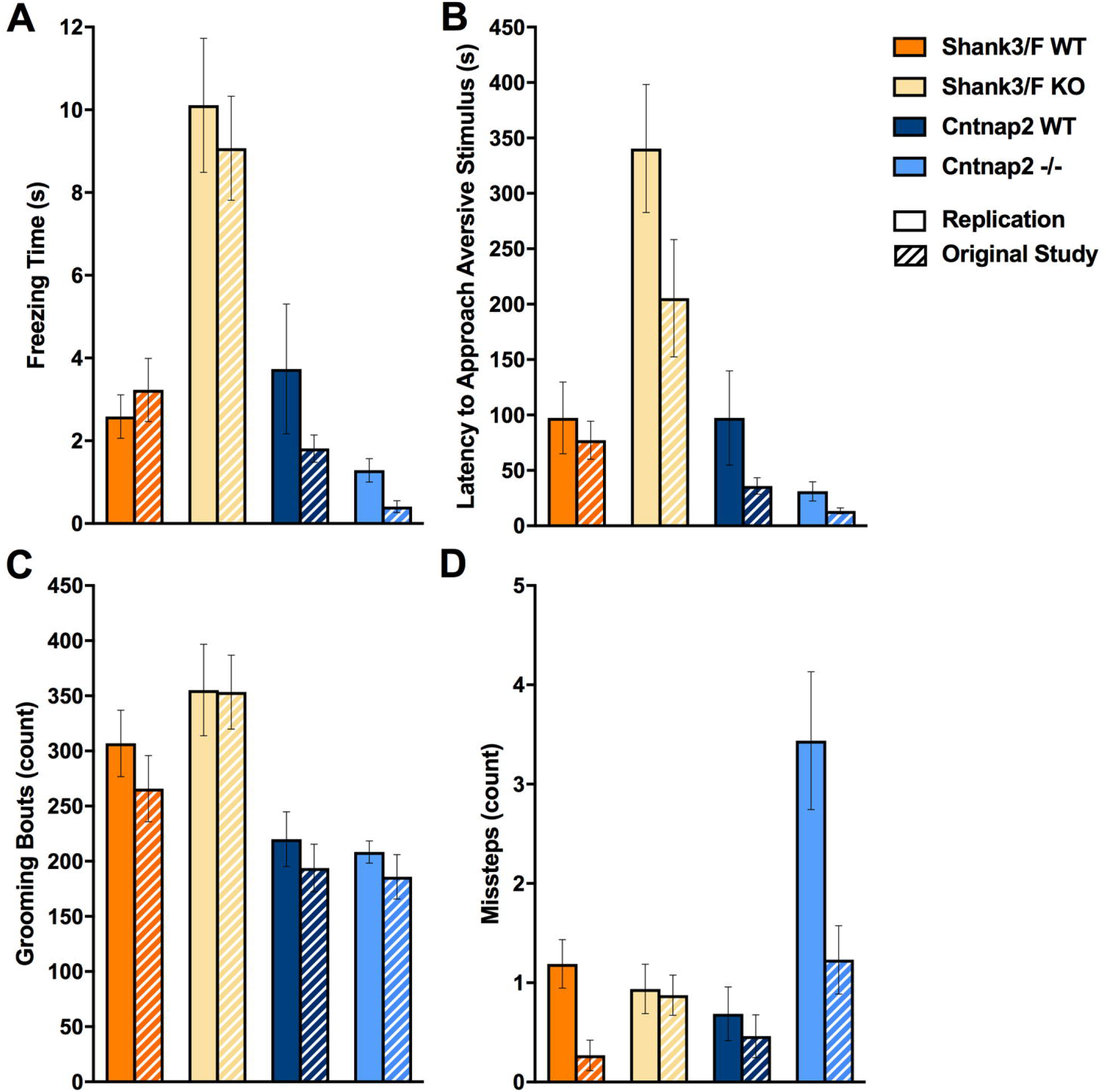
Additional SmartCube^®^ top features showed differences across the models that were consistent across both studies. The top features that separate the mutant and control groups are plotted for both models and for the replication and original (hashed bars) study. *Shank3/F* KO mice showed (or tended to show) increased freezing (A), latency to approach an aversive stimulus (B), and grooming (C) compared to WT littermates, whereas the *Cntnap2* -/- mice showed again no difference or decreases in measures compared to WT littermates. Misstepping (D) was the only top feature that showed inconsistent results across models and studies.

The top features for the replication and historical studies are shown in Figure 3 and Figure 4 (S1 Table). It is clear that the two model systems show opposite differences or trends that are surprisingly consistent between the historical and replication studies. For example, whereas the *Shank3/F* KO mice showed increased freezing in both studies, the *Cntnap2*-/- mice exhibited less freezing. The WT groups, which are both C57/B6 littermates, show similar freezing levels.

Figure 5 shows an analysis of all five model systems together. In the top panel (see Fig A) we combined the replicas for the *Shank3/F* and *Cntnap2* model systems, whereas in the bottom panel (see Fig 5B) the replicas are shown separated. To run this analysis, we trained the classifiers on all model systems at the same time, aiming at separating all of them from each other as much as possible. Surprisingly, models landed heavily on a line, with the combined WT group in the center. Thus, the *Shank3* model systems were on one side of the combined WT group, and the *Cntnap2* on the other side. The 16p11.2 and *Cacna1c* model systems did not show a strong signature. When replicas were run separately they landed very much on top of each other and again on a line, suggesting good replication and opposite phenotypes for the *Shank3/F* and *Cntnap2* model systems.

**Fig 5.**
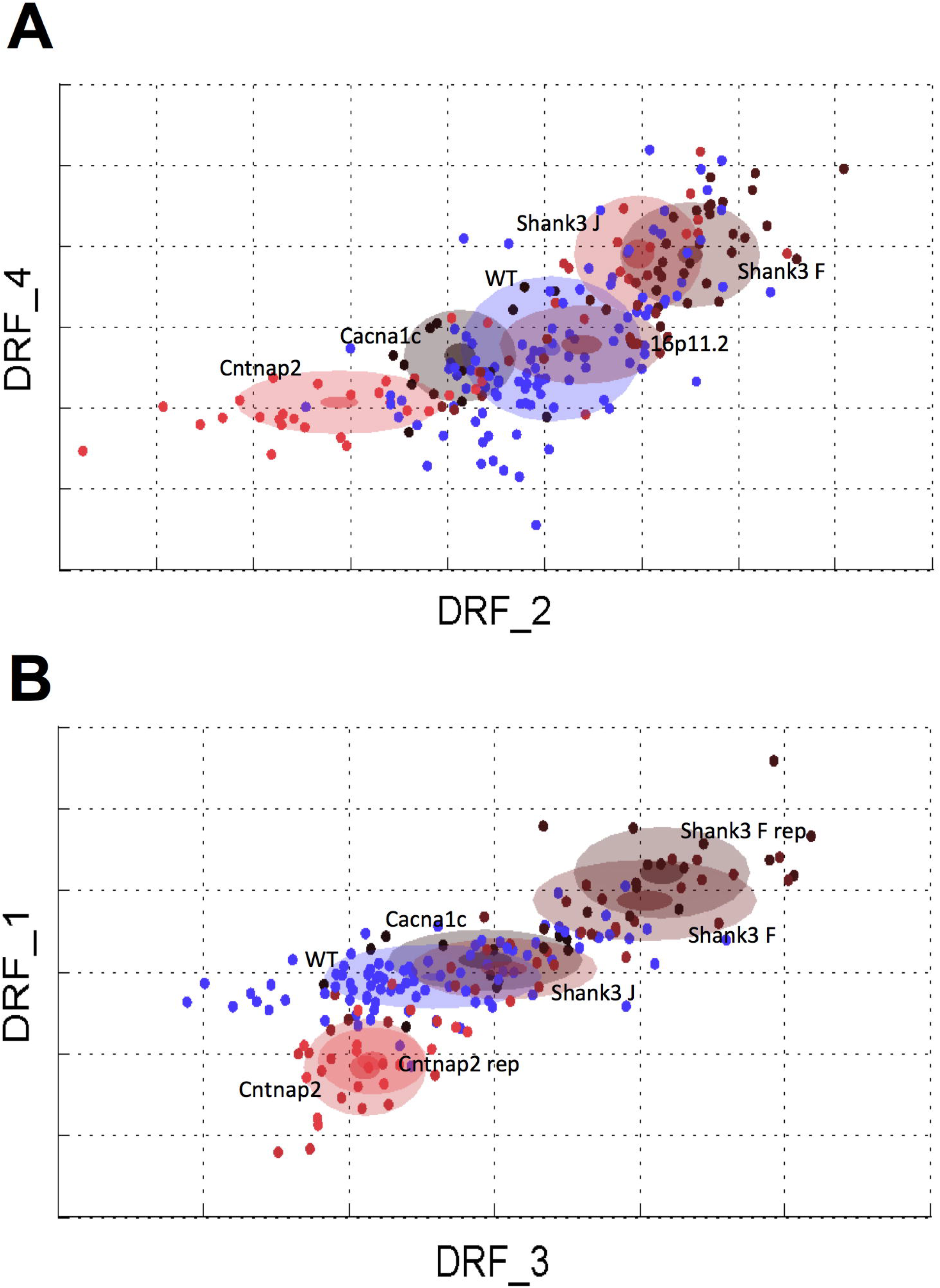
“Cloud” analysis across ASD mouse models. A: All five ASD mouse models plotted in the same multidimensional space with mutant replicas and WT littermates pooled. B: The four mouse models on C57/B6 genetic background plotted in the same multidimensional space with mutant replicas separated and WT littermates pooled. N=16 mice per genotype/line (replication), n=13-16 mice per genotype/line (original studies); dots are individual animals.

##### Body Weight

All groups significantly gained weight with age during the study (Fig 6; S2 Table). The *Shank3/F* KO mice tended to be slightly heavier than WT littermates but this did not reach significance in the replication, unlike in the original study (at P90). A non-significant trend for decreased weight was also observed in the *Cntnap2* -/- mice, resembling the significant differences found in the first study (at P90). It should be noted, however, that in original study two cohorts of mice were needed to run all tests in the comprehensive phenotypic screen and therefore the sample size is double in size (n=28-32 per group) compared to the replication (n=16 per group).

**Fig 6.**
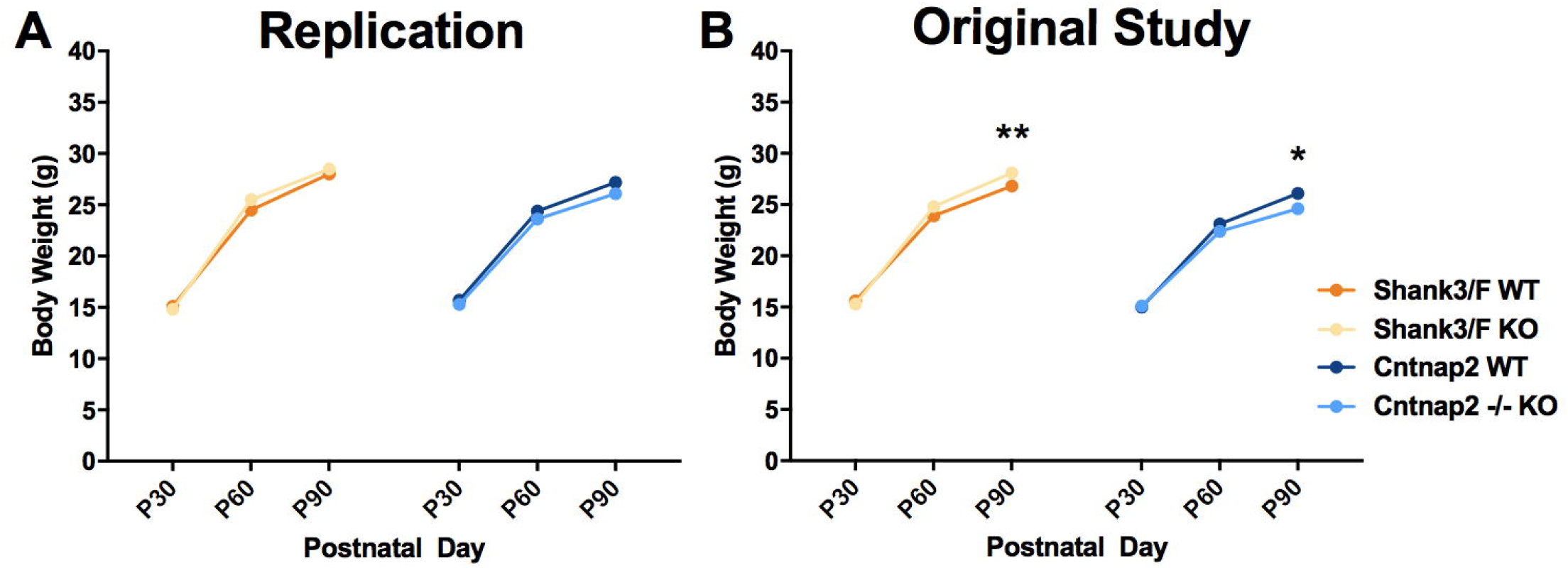
Body weight. No differences in body weight were found between mutant and wild type mice for either model in the replication study. Note errors bars are hidden by the graphing symbols. Data shown are means ± SEM; n=16 mice per genotype/line (replication); n=28-32 mice per genotype/line (original study); \**p*< 0.05; \*\**p*< 0.01.

##### Reciprocal Social Interaction

To assess social behavior, we paired genotype- and age-matched mice and allowed them to interact freely for 10 minutes. The replication results matched very closely the original study data (S3 Table). *Shank3/F* mutant pairs were or tended to be closer to each other, and remained so for longer, than the corresponding WT pairs (Fig 7A and 7B). However, they also followed each other less and showed less locomotion (Fig 7C and 7D), suggesting an effect of hypoactivity on social behavior. Indeed, *Shank3/F* mutant pairs interacted less frequently with each other but when they did so, they interacted for longer than WT controls (Fig 8A and Fig 8B), again, consistently with a hypoactive profile.

**Fig 7.**
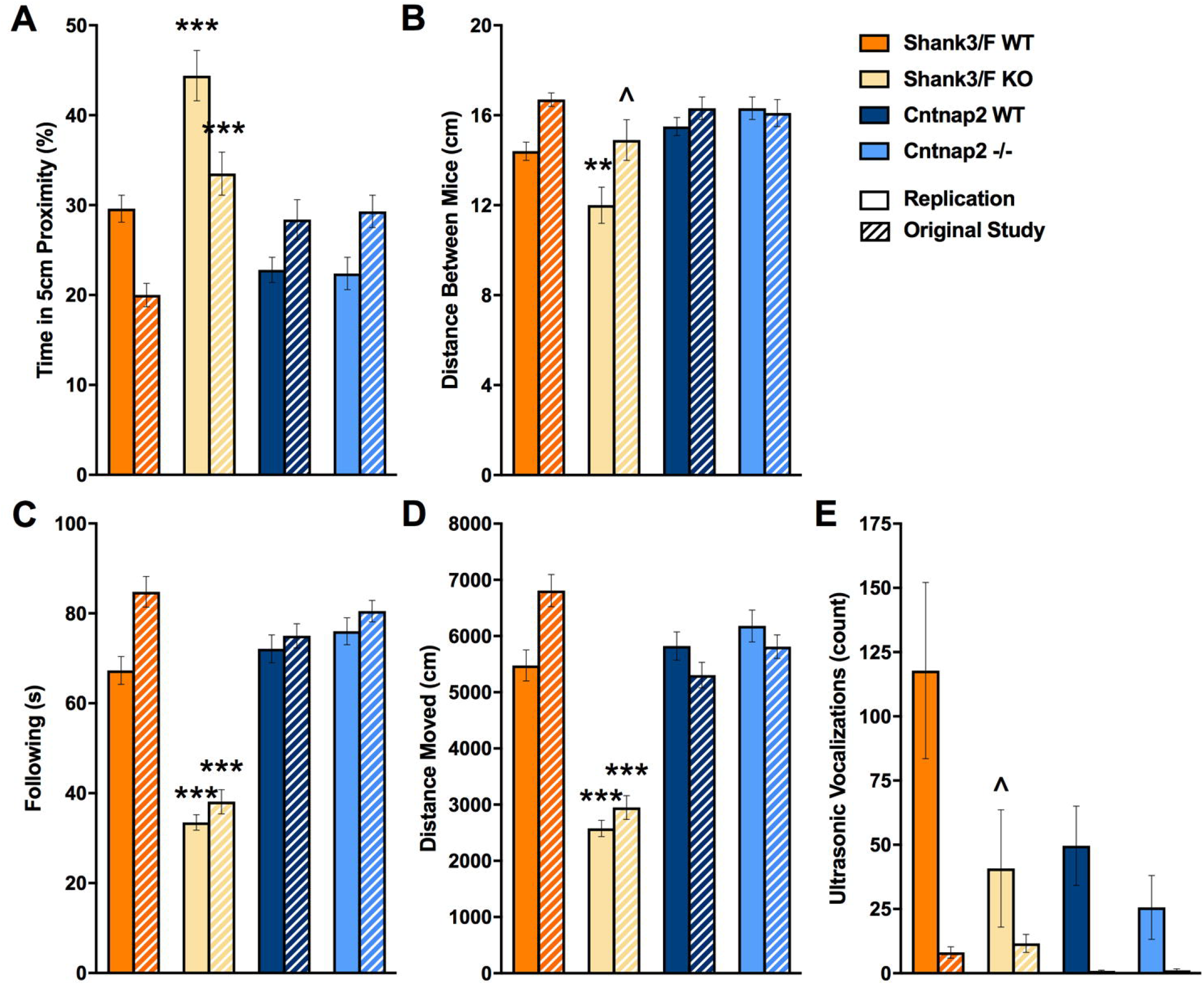
Proximity, activity, and vocalizations emitted during reciprocal social interaction. Measures are shown for the replication and original (hashed bars) study. A: *Shank3/F* KO pairs spent more time in close proximity to each other than the corresponding WT pairs in both the original and replication study, whereas no phenotypic differences were seen in the *Cntnap2* model across both studies. B: *Shank3/F* KO pairs were in closer proximity to each other than the corresponding WT pairs in the replication study, whereas the difference was only marginal in the original study. There were no differences in the *Cntnap2* model across both studies. C: The time following each other was decreased in the *Shank3/F* KO as compared to the WT mice across both studies. The *Cntnap2* model did not show a phenotypic difference in either study for this measure. D: Distance moved showed a very similar pattern to following. E: *Shank3/F* KO mice tended to emit fewer vocalizations than WT littermates in the replication but not in the original study, whereas no significant differences were observed in the *Cntnap2* model in either study. Data shown are means ± SEM; n=16 mice per genotype/line (replication), n=14-16 mice per genotype/line (original study). (^*p*< 0.08, \*\**p*< 0.01, \*\*\**p*< 0.001).

**Fig 8.**
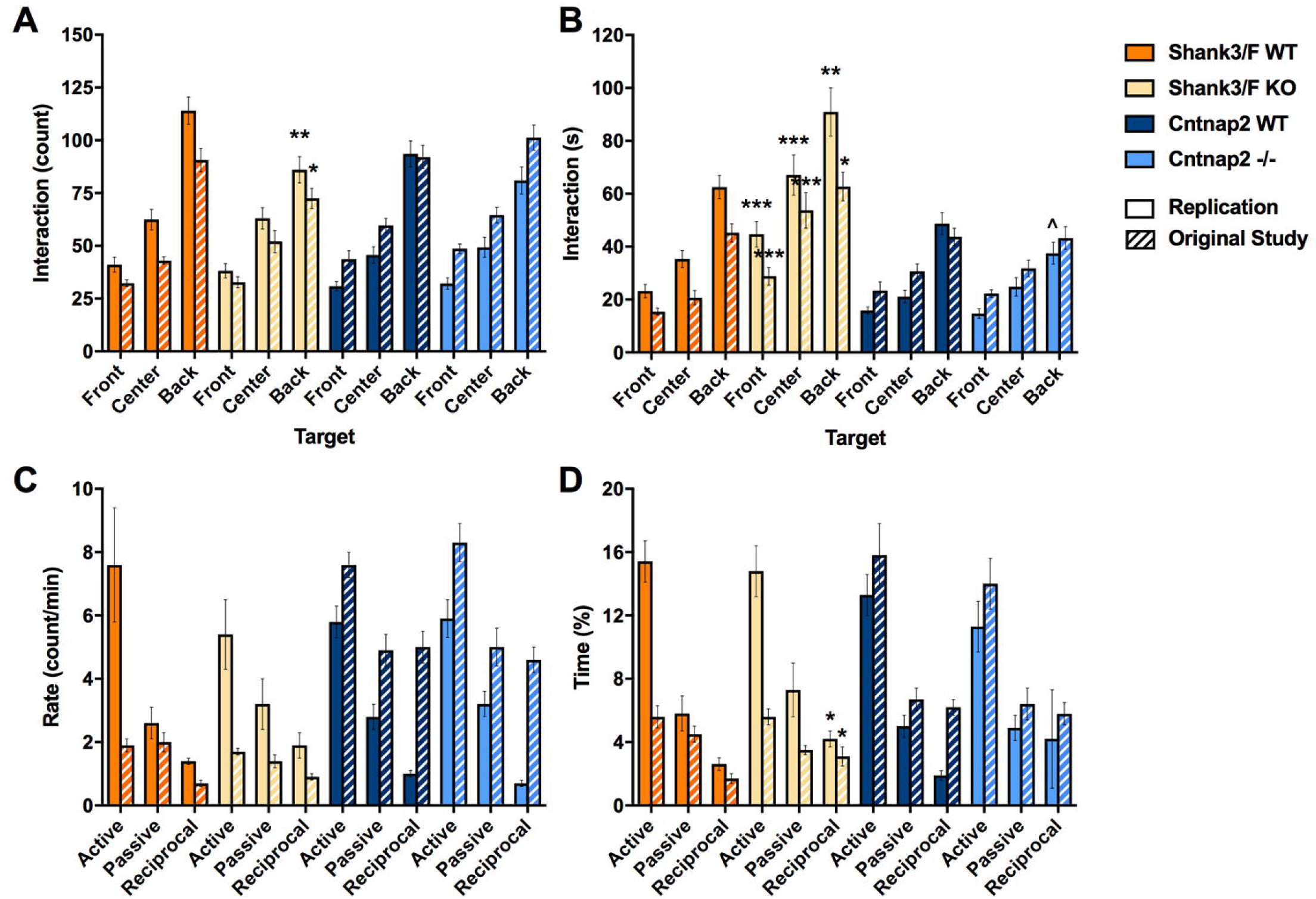
Social interactions. Measures are shown for the replication and original (hashed bars) study. A: *Shank3/F* KO pairs showed fewer interactions (approaches towards each other’s back) than the corresponding WT pairs, and no differences were observed for *Cntnap2* pairs. These results were identical in both the original and the replication studies. B: The time interacting, however, showed opposite patterns in the *Shank3/F* mice, with the KO pairs interacting for a shorter duration than the WT pairs. Again no differences were observed in the *Cntnap2* pairs. Once again, the results were identical in the original and replication studies. C: Hand scoring did not capture differences in the frequency of interaction in either study (“Rate”), for either model. D: The percent time interacting, however, showed increased reciprocal interactions in the *Shank3/F* KO mice compared to WT littermates, and, again, no differences in the *Cntnap2* model in both studies. Data shown are means ± SEM. These results were identical across the two studies. (^*p*< 0.08, \**p*< 0.05, \*\**p*< 0.01, \*\*\**p*< 0.001).

In addition to automated scoring, social interactions were hand-scored to isolate behaviors driven by the *subject* mouse (which was introduced to the testing chamber five minutes before the *stimulus* mouse). We found no differences in the rates of active (subject investigating the stimulus mouse), passive (stimulus investigating the subject mouse), or reciprocal (subject and stimulus mouse investigating each other) interactions (Fig 8C). However, increased time spent in reciprocal investigation was observed for *Shank3/F* KO mice across both studies (Fig 8D). The only measure not entirely confounded by activity in this test was ultrasonic vocalizations (Fig 7E). Although vocalizations were too infrequent to analyze in the original study, a trend of decreased vocalizations was observed in *Shank3/F* KO mice. No notable differences were observed between *Cntnap2* -/- and WT mice across either study with slight trends to show decreased vocalizations not reaching statistical differences (Fig. 7E).

##### Urine-Exposure Open Field

To assess social behavior in a different way, potentially less confounded by the behavior of companion mice and the levels of motor activity, we used an open field test in which male mice are exposed to urine of a female in estrous for five minutes after a one-hour baseline session (for all results see S4 Table). Whereas sex-naïve mice readily scent-mark (Novotny, Harvey et al. 1990, Lehmann, Geddes et al. 2013), ultrasonic vocalizations require social experience. Therefore, we exposed all males to females one week prior to the open field test.

*Shank3/F* KO mice were again hypoactive in this test as compared to the WT pairs, showing less locomotion around the chamber and in the center during both the baseline and urine exposure sessions in the replication study (Fig 9). This pattern did not reach statistical significance during the baseline session in the original study (Fig. 9A and 9C), but showed the same difference for the urine test phase (Fig. 9B and 9D). *Shank3/F* KO mice additionally showed less time in the center of the open field as compared to the WT pairs in both the baseline and the urine-exposure session and fewer ultrasonic vocalizations (Fig 10A and Fig 10B). These results were identical in the original and replication studies. Although there were no differences in overall chamber scent marking, it was significantly decreased in the center after urine exposure in the replication study and showed a similar trend in the original study, suggesting that this behavior may be indicative of a social deficit (Fig 10C and Fig 10D).

**Fig 9.**
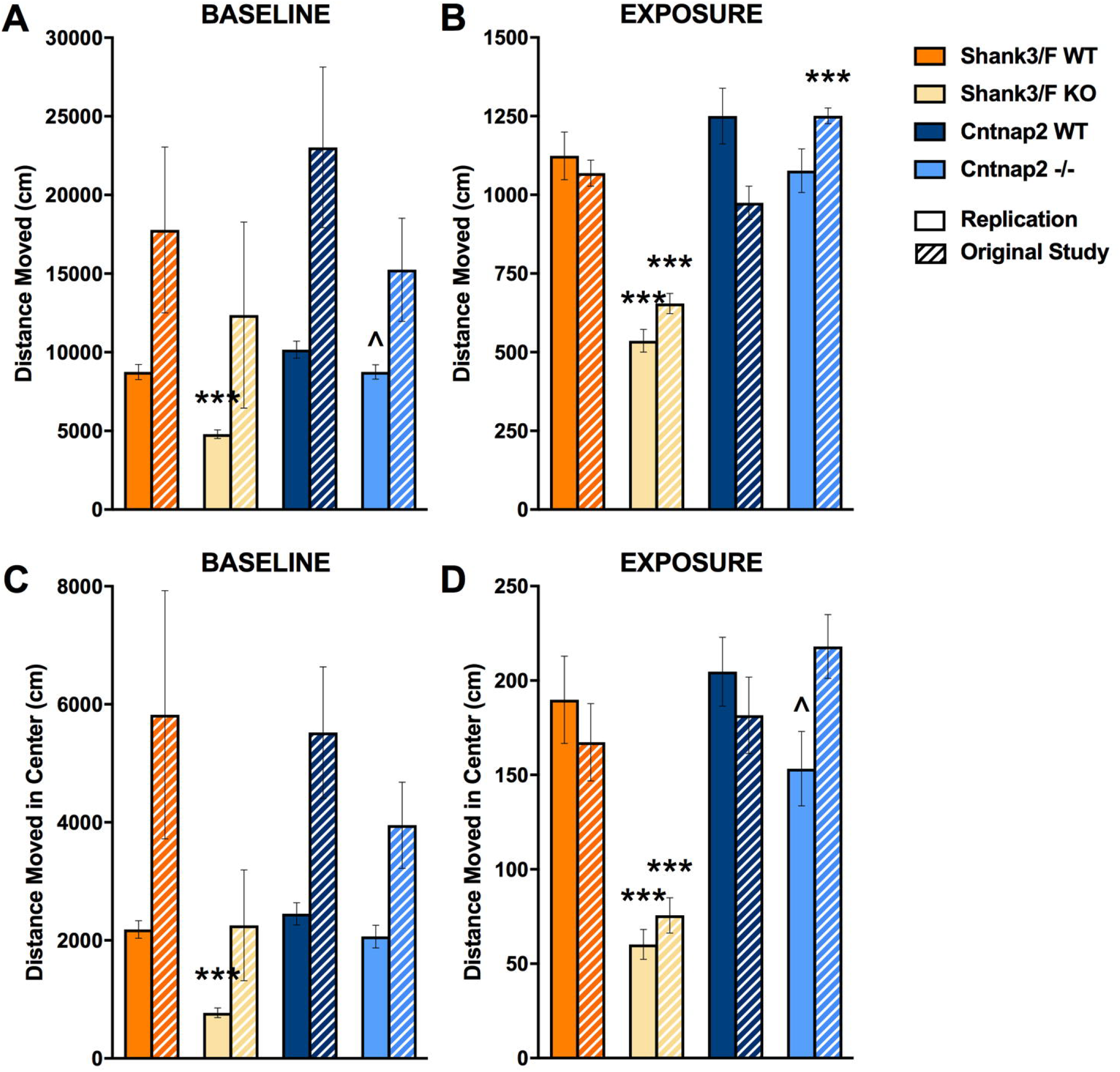
Open field activity during the baseline and urine-exposure sessions. Measures are shown for the replication and original (hashed bars) studies. A: During the baseline session, *Shank3/F* KO mice moved less and *Cntnap2* -/- tended to move less around the chamber than WT littermates during the replication but not in the original study. B: Shank3/F KO mice moved around the chamber less than WT mice during the exposure session in both studies while *Cntnap2* -/- mice moved around the chamber more than WT mice in the original study only. C: *Shank3/F* KO mice moved less in the center than WT mice during the baseline session in the replication but not the original study, whereas there were no differences between KO and WT in the *Cntnap2* model. D: *Shank3/F* KO mice moved less in the center than WT mice during the urine-exposure session in both studies while *Cntnap2* -/- mice only tended to move less in the center during urine exposure in the replication study. Data shown are means ± SEM; n=16 mice per genotype/line (replication), n=15-16 mice per genotype/line (original study). (^*p*< 0.07, \*\*\**p*< 0.001).

**Fig 10.**
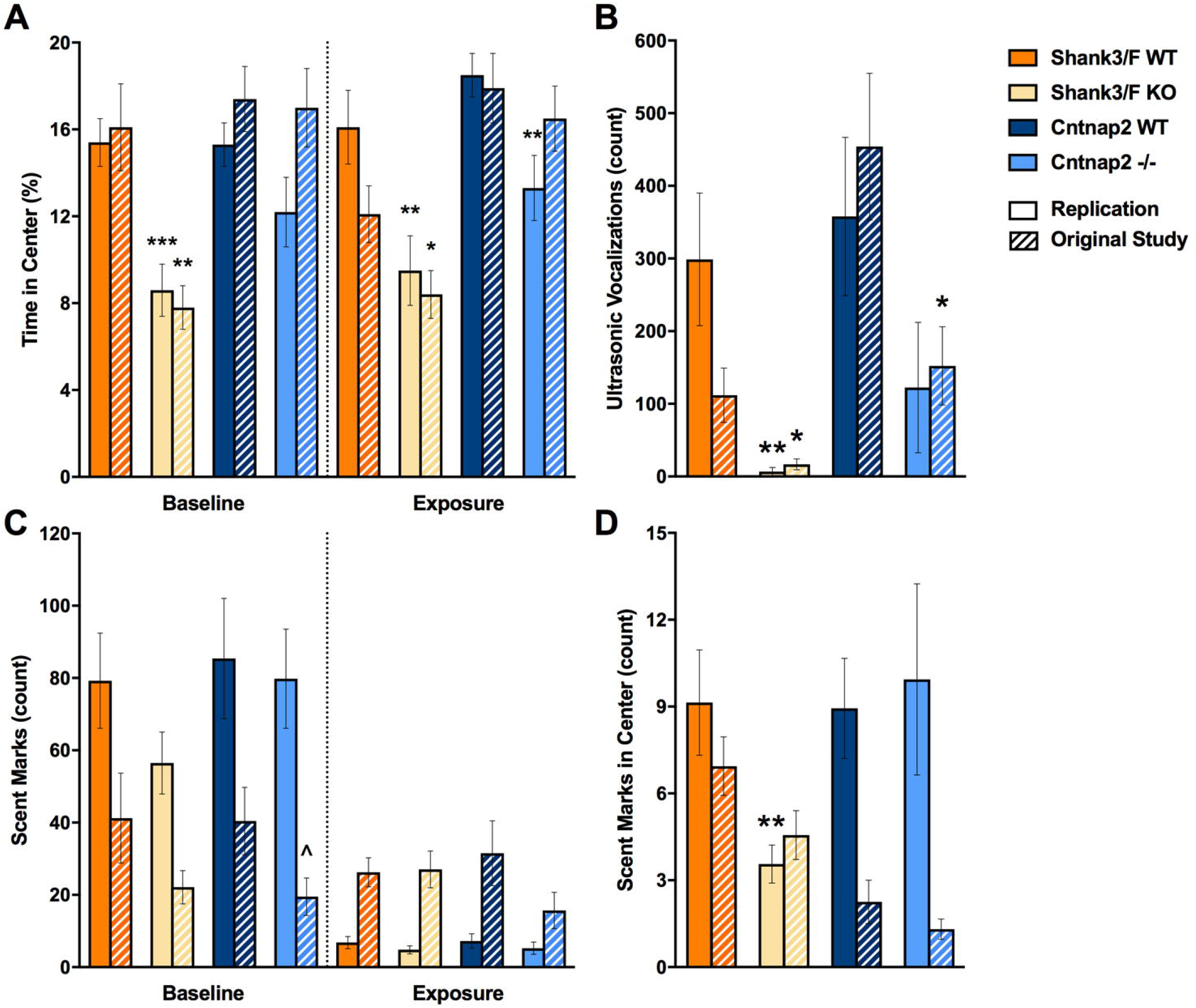
Time in the center, ultrasonic vocalizations, and scent marking in the urine open field test. Measures are shown for the replication and original (hashed bars) study. A: Shank3/F KO mice spent less time in the center than WT mice during both testing sessions across both studies, while *Cntnap2* -/- mice only spent less time in the center during urine exposure compared to WT mice in the replication study. B: *Shank3/F* KO male mice vocalized less than WT mice in both studies, whereas *Cntnap2* -/- mice vocalized less than WT mice only in the original study. C: KO mice from both models in both studies scent marked similarly to WT mice during both baseline and urine exposure sessions. D: *Shank3/F* KO male mice scent marked less than WT mice in the center during the urine-exposure session in the replication study only while there were no differences between KO and WT in the *Cntnap2* model. Data shown are means ± SEM. (^*p*< 0.07, \**p*< 0.05, \*\**p*< 0.01, \*\*\**p*< 0.001).

*Cntnap2* -/- mice did not exhibit a clear hyperactive phenotype in this test. During the baseline session, they tended to locomote less around the chamber as compared to the WT pairs, but this was only a statistical trend in the replication study (Fig 9A and Fig 9C). Behavior during the exposure session was quite inconsistent; with *Cntnap2* -/- mice showing more locomotion around the chamber as compared to the WT pairs in the original study and slightly less locomotion in the center in the replication study (Fig 9B and Fig 9D). These mutant mice also showed less time in the center, but only during the urine exposure session and only in the replication study (Fig 10A). In the historical data, there were no differences or trends. Ultrasonic vocalizations were reduced in the original study but a trend in the same direction did not reach significance in the present study (Fig 10B). Overall scent marking was slightly reduced in the original study but otherwise did not show any differences (Fig 10C and Fig 10D).

The *Shank3/F* mice, therefore, showed an anxiety-like phenotype, and reduced social ultrasonic response as compared to the WT pairs and scent marking near to a female stimulus, whereas results in the *Cntnap2* model system were not consistent.

#### Interlaboratory Replication and Assessment of Convergent Validity

##### Open Field – Standard Test of General Activity

To provide a standard measure and characterization of activity we ran a 60-min open field test (for all results, see S5 Table). Whereas in our previous studies we assessed the development of motor function in juvenile mice, we focused here on assessment of the young adults (∼P75), to provide a separated assessment of activity at the age when we assessed social behavior, since for some of these tests activity may be a confounding factor. We cannot, therefore, assess direct replicability of activity test *per se*, rather we extend measures of activity to an older age. Distance covered was reduced overall in both model systems, as was stereotypic behavior (Fig 11A and Fig 11C). Rearing (Fig 11B), and all measures in the arena center (Fig. 12: A - time, B - distance moved, C-rearing, D - stereotypic counts) were only decreased in the *Shank3/F* KO mice, relative to the WT control mice.

**Fig 11.**
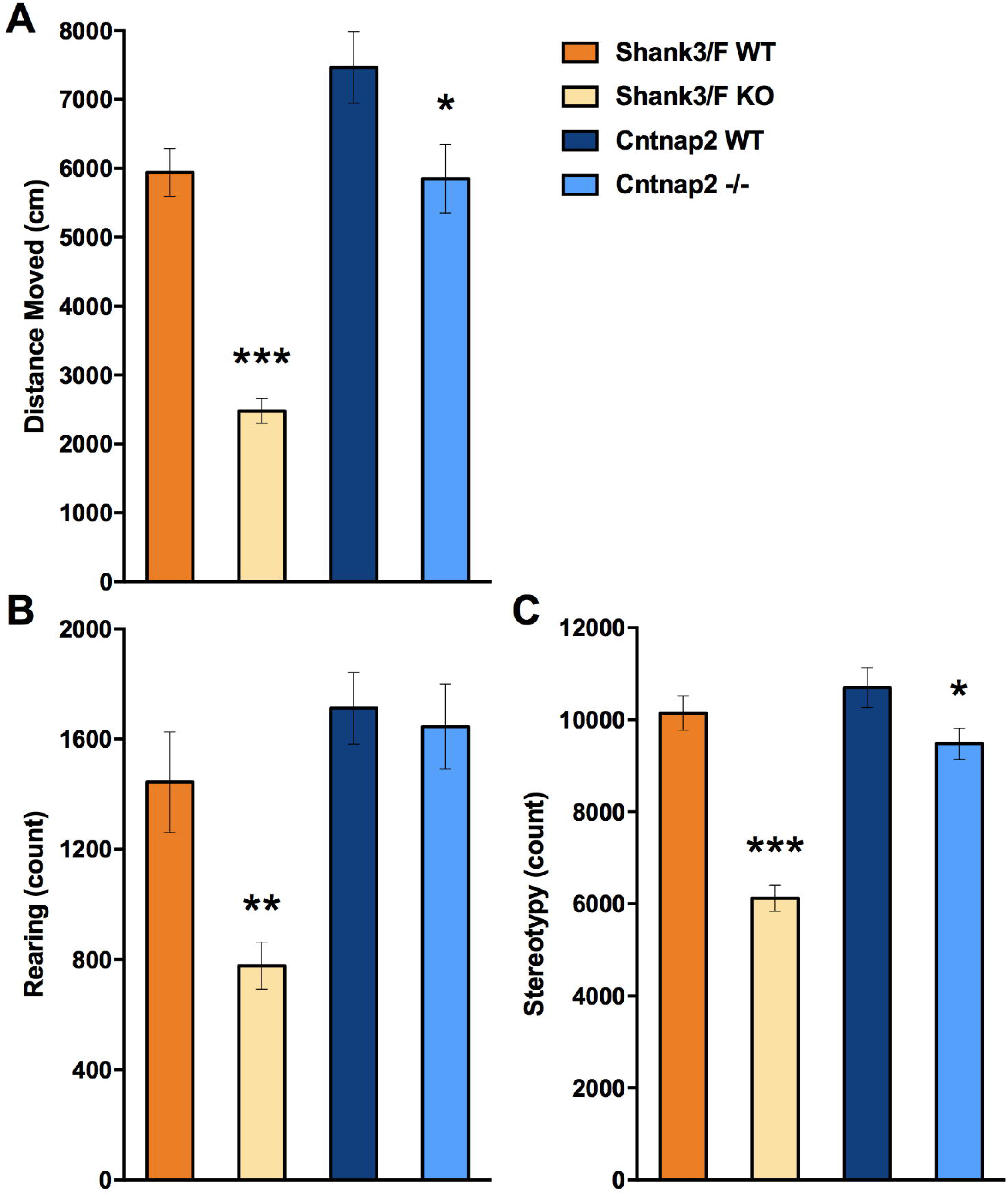
Overall behavior in the standard open field test. *Shank3/F* KO mice moved (A), reared (B) and showed less stereotypic behavior (C) than WT littermates. *Cntnap2* -/- mice moved (A) and showed less stereotypic behavior (C) than WT littermates, but did not differ in rearing frequency (B). Data shown are means ± SEM; n=16 mice per genotype/line. (\**p*< 0.05, \*\**p*< 0.01, \*\*\**p*< 0.001).

**Fig 12.**
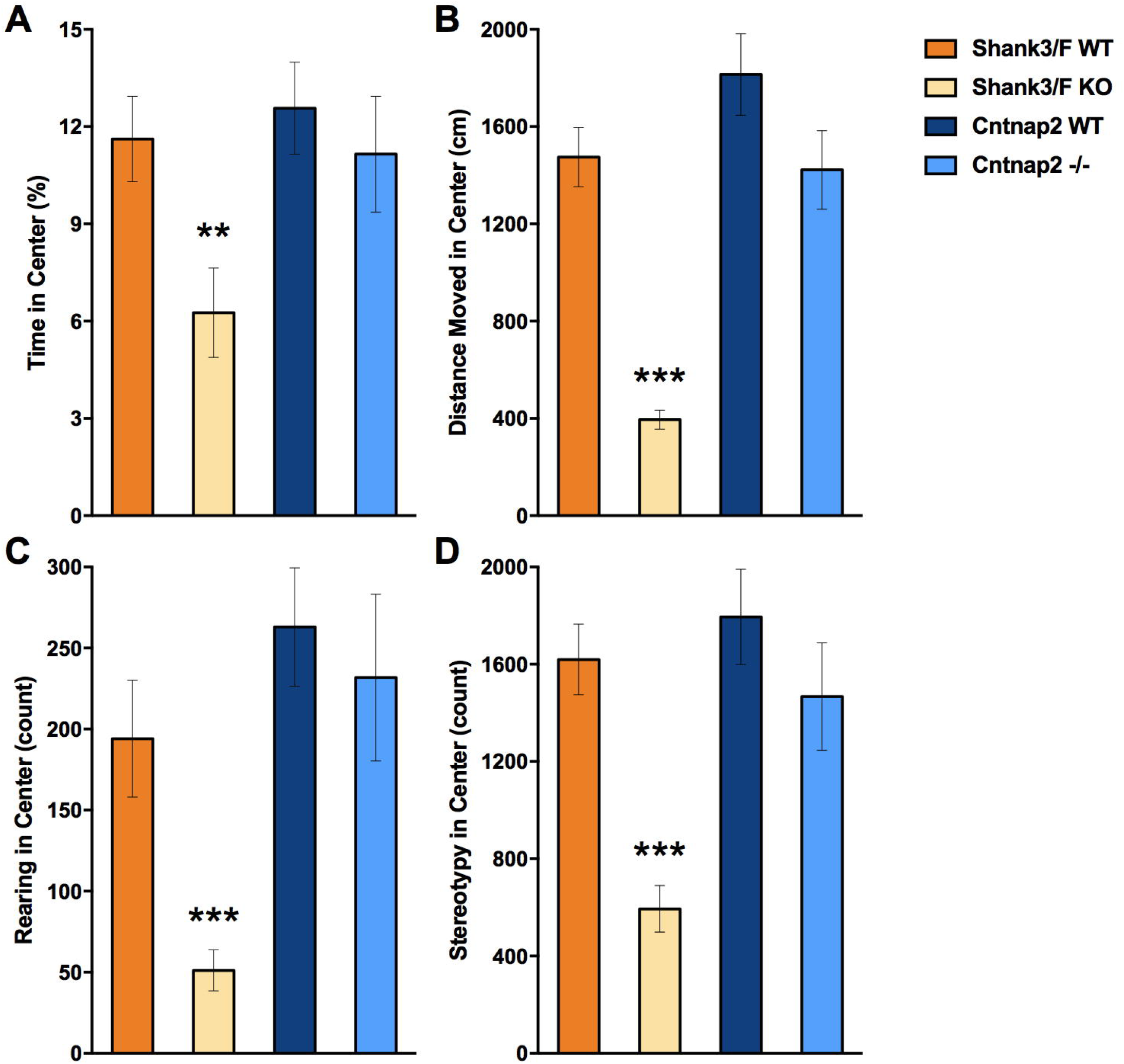
Behavior in the center during the standard open field test. *Shank3/F* KO mice spent less time (A), moved (B), reared (C), and showed less stereotypic behavior (D) in the center of the open field as compared to WT control mice. *Cntnap2* -/- mice did not differ from WT control mice on any measure in the center of the open field. Data shown are means ± SEM. (\*\**p*< 0.01; \*\*\**p*< 0.001).

##### Grooming – Test of Repetitive/Anxiety-like Behavior

Following the Peça et al. (2011) report of increased time grooming in the *Shank3/F* KO mice, we assessed grooming in long sessions of 120 min (for all results see S6 Table). Whereas both model systems showed increased grooming frequency (Fig 13B) compared to WT littermates, the increase in time only reached significance for the *Cntnap2* -/-model system (Fig 13A).

**Fig 13.**
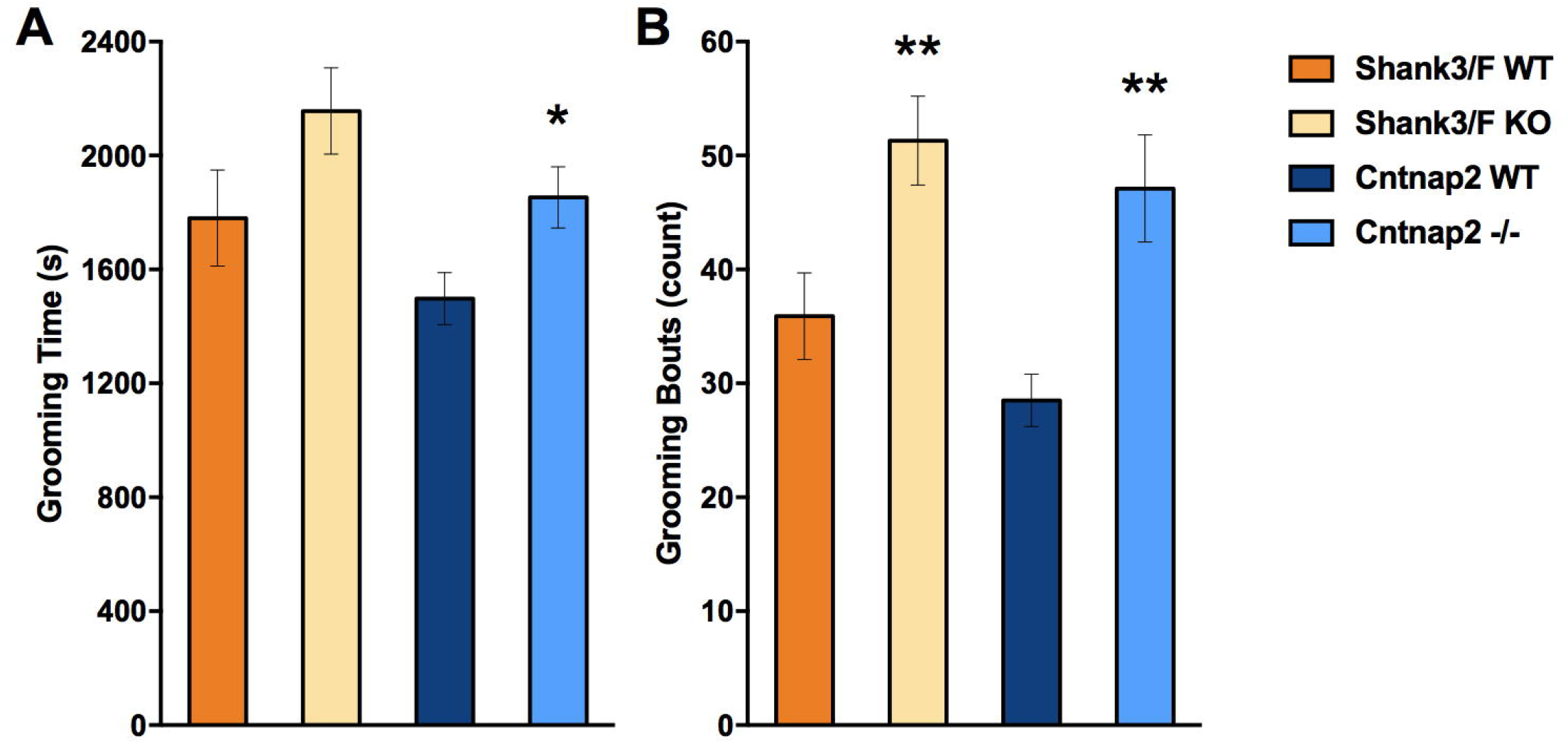
Grooming. *Cntnap2* -/- mice spent more time (A) and showed more bouts (B) of grooming than WT littermates, whereas *Shank3/F* KO mice just showed more bouts (B) of grooming. Data shown are means ± SEM; n=15-16 mice per genotype/line. (\**p*< 0.05; \*\**p*< 0.01).

### 5.4. Discussion

#### Findings Across the Three Studies

Animal model systems are needed in neuroscience and drug discovery to better understand the fundamental pathology and pathogenesis of disease, identify and validate drug targets, and screen potential therapeutics. The use of animal models, however, presents many challenges. In ASD, for instance, there are many models based on different genetic findings, from gene mutations to copy number variation, for which phenotyping efforts typically concentrate on three main domains of ASD, namely, repetitive behavior, communication, and social deficits. More often than not, researchers assume implicit homologies of the substrates underlying the symptoms in humans and those in rodents, using a simple face value similitude principle. For instance, social behavior in mice is used to model social deficits in children with ASD, and ultrasonic vocalizations to model communication deficits. Apart from the implications of such homology assumptions, which can and should be challenged, this simple approach implies that all models of ASD will present with similar behavioral patterns. Our project aimed at comparing five different ASD models comprising a large phenotypic effort (Brunner, Kabitzke et al. 2015, Kabitzke, Brunner et al. 2018) and examining these implicit assumptions. In addition, we studied the robustness and reproducibility of the results. Therefore, we present here a brief summary of all our previous results, replications of several new results using the Shank3/F and Cntnap2 ASD models, and conclusions from machine learning based tests that identified putative common behavioral features across the five model systems investigated.

##### Development

In terms of development, all model systems showed a rather normal and robust progression (see Table 2). The only exception was the 16p11.2 HET mice, which were smaller than the corresponding WT mice. This is consistent with the finding that this model system is lethal in the C57 genetic background (Horev, Ellegood et al. 2011), suggesting some serious early pre- or postnatal issues. *Shank3/F* KO mice were very slightly heavier, consistent with previous publications (Wang, McCoy et al. 2011), although the replication study showed just a non-significant trend, whereas the *Cntnap2* mutant mice were very slightly underweight.

**Table 2.**
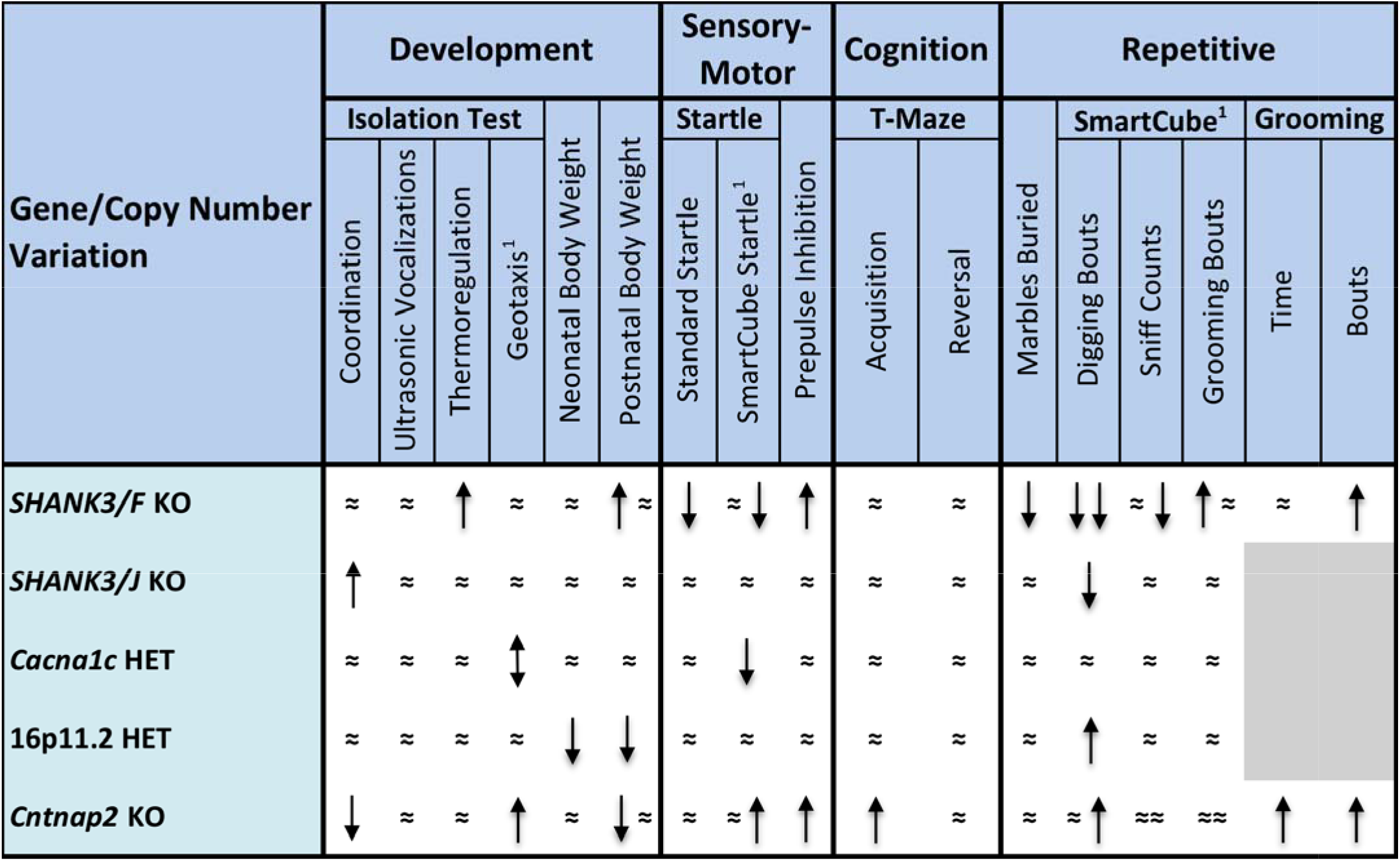
Summary of developmental, sensory-motor, cognitive, and repetitive behavioral results across the ASD animal models studied. Directions of effects are shown comparing each model system to their corresponding WT controls (upward arrow: increased, downward arrow: decreased, ≈: no difference). Two symbols occupying a single cell reflect both the original study and replication results; ^1^ quantitative measure (see the description of test assessment in Methods); grey field indicates no data collected for that model.

##### Neonatal Ultrasonic Vocalizations

The measure of USV was quite variable suggesting that much larger sample sizes may be required to find robust differences (see Table 2). Alternatively, longer isolation sessions can be used, although the risk of causing unknown long term changes needs to be examined carefully if the subjects are to be reused later in development (e.g., Scattoni, Ricceri et al. 2011).

##### Startle Response

*Shank3/F* KO mice showed lower startle levels than the corresponding WT mice in two different tests (ASR and SmartCube^®^ tests), whereas the *Cancnac1* mice showed a similar pattern of reduced startle response compared to WT mice in the SmartCube^®^ test only (see Table 2). These findings are of interest given that *Shank3* has been suggested to directly contribute to Rett pathology, and mouse model systems of Rett show a robust startle deficit (Waga, Asano et al. 2014). It is tempting to speculate that the link between the startle deficit in the *Cacna1c* and *Mecp2* mouse model systems originates from BDNF modulation of hippocampal neurogenesis, which is affected in both model systems (Larimore, Chapleau et al. 2009, Lee, Hill et al. 2016). If this hypothesis is supported by other evidence (such as replication of the deficit in a standard test) it would be interesting to use startle as a way to screen drugs for ASD, as is currently done in Rett syndrome (Brunner & Kabitzke, unpublished drug screen platform), given the good homology between the startle circuitry between rodents and humans. Different SHANK3 isomorphs could lead to different phenotypes, and this may be the reason that *Shank3* from Jiang’s lab did not show a startle deficit (Zhou, Kaiser et al. 2016). The *Cntnap2* mutant mice showed an increase in startle in SmartCube^®^ as compared to the WT controls but not in the standard test, a finding that would also require replication. Interestingly, PPI was increased in both the *Shank3/F* and *Cntnap2* mutant mice, a finding opposite to what one would expect for schizophrenia, but consistent with some findings in ASD (Madsen, Bilenberg et al. 2014).

##### Repetitive Behavior

We used several tests to assess repetitive behavior (see Table 2). An interesting pattern emerged in the *Shank3/F* mice, which showed decreased marble burying, digging, and sniffing compared to WT littermates but increased grooming frequency, consistent with previous reports (Wang, McCoy et al. 2011). The *Cntnap2* mice also showed increased grooming compared to WT littermates, consistent with previous findings, but no changes in marbles buried (Bader, Faizi et al. 2011). Few tests in this battery showed similar patterns between the *Shank3/F* and *Cntnap2* model systems, the two models out of the five studied that presented the more robust phenotypes, and thus it is of particular interest to confirm perseverative grooming. This is another simple endpoint measure with good translation potential, as the circuitry involved is very conserved across species (Kalueff, Stewart et al. 2016). Opposite patterns in these tests suggest differing neuroanatomical and neurotransmitter support, arguing against the interchangeable use of these tests for assessment of perseveration.

##### Locomotor Activity

Consistent with previous reports (Bader, Faizi et al. 2011, Wang, McCoy et al. 2011), several model systems showed a decrease in locomotor activity compared to their WT controls (see Table 3). Whereas the two *Shank3* and the *Cacna1c* model systems were consistently hypoactive across different tests as compared to their respective WT mice, the *Cntnap2* mice were hypoactive in the open field, did not show strong differences in the urine open field or marble burying, but was nevertheless hyperactive in the SmartCube^®^ test. This SmartCube^®^ test was designed to provide strong stimulation and cause strong reactions (such as a defensive burying response to an electrical probe, and a startle response to tactile stimulation), and thus it is possible that the pattern of activity in the *Cantnap2* mice reflect environmental reactivity, whereas the *Shank3/F* mice show a low endogenous level of activity, regardless of environmental stimulation. Our informatics analysis suggested that, at least in the SmartCube^®^ test, there is a signature continuum, with the model systems lining up in the following order: *Shank3/F* -> *Shank3/J* -> *Cacna1c* -> all WT & 16p11.2 -> *Cntnap2*. Interestingly, independent of the distance covered during locomotion, most model systems moved faster in the NeuroCube^®^ test. SmartCube^®^ did reproduce for a second time the published findings of reduced rearing in *Shank3/F* KO mice compared to WT mice (Peça, Feliciano et al. 2011), consistent with its overall hypoactive pattern.

**Table 3.**
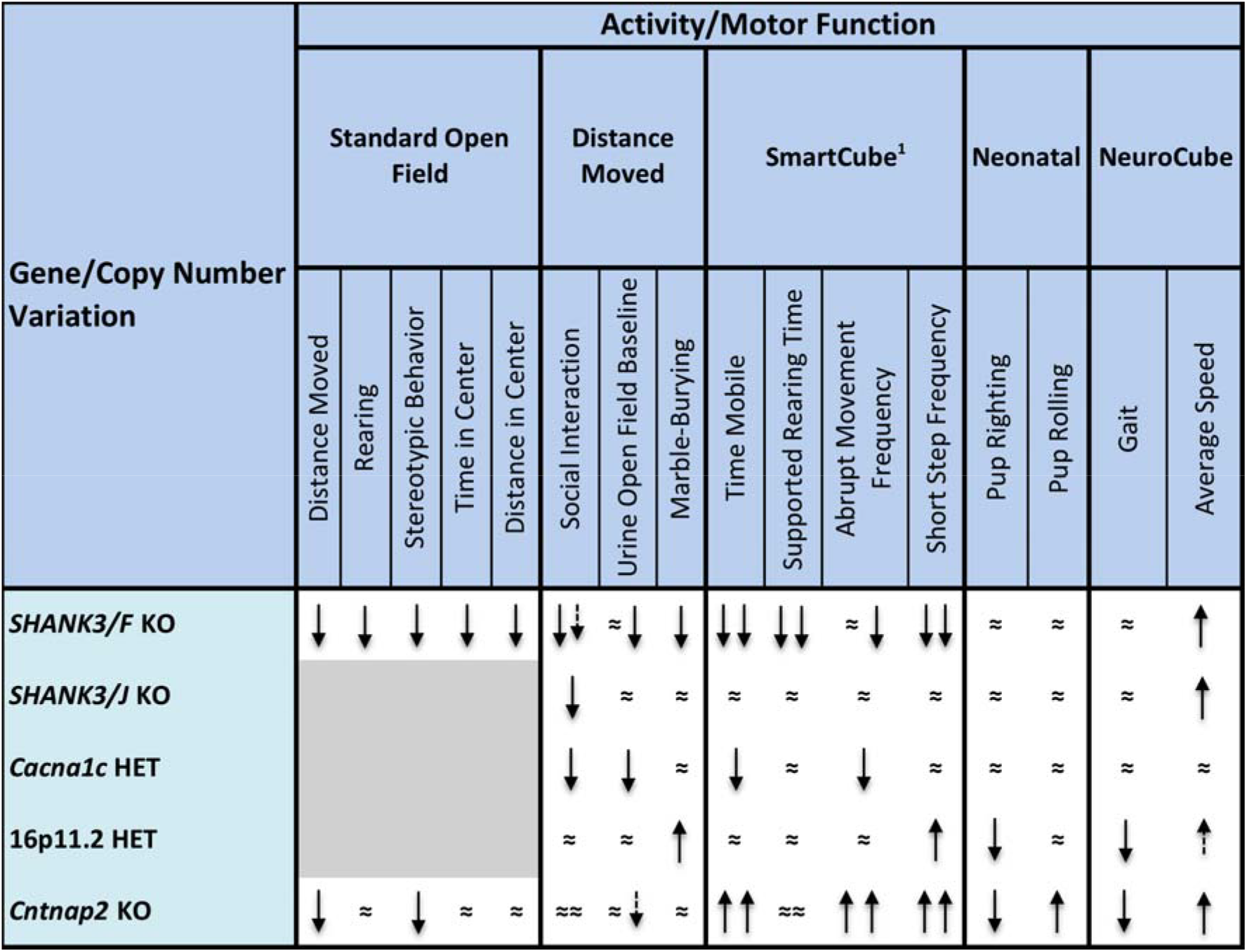
Summary of activity/motor function results across the ASD animal models studied. Directions of effects are shown comparing each model system to their corresponding WT controls (upward arrow: increased, downward arrow: decreased, ≈: no difference, dashed shorter arrow: non-significant trend). Two symbols occupying a single cell reflect both the original study and replication results; ^1^ quantitative measure (see the description of test assessment in Methods); grey field indicates no data collected for that model.

##### Social Behavior

We used three different tests to analyze behavior in a social context: the three chamber, reciprocal interaction, and urine-open field tests (see Table 4). We previously reported that *Shank3/F* KO mice showed no preference for the social stimuli in the three-chamber test but that perhaps genotypic differences did not reach significance due to the high variability in the WT group (Kabitzke, Brunner et al. 2018). The lack of significance in the other model systems, however, did not seem confounded by any other factor and probably reflects no deficit (or undetectable deficits) in our experimental setup. In the reciprocal social interaction test, we found that mice that were hypoactive (*Shank3/F, Shank/J*, and *Cntnap2*) follow each other less, but interact for longer periods of time. This strongly suggests a confounding effect of activity, in particular for the *Shank3/F* and *Cacna1c* models, as they were hypoactive in different tests of motor activity. The 16p11.2 mice also showed closer proximity and more time spent interacting, clearly not a social “deficit”. Finally, the *Cntnap2* mice were not significantly different from the corresponding WT group across any of the reciprocal social interaction measures. Social behavior in rodents, and associated learning and recall processes, primarily depends on olfactory function, whereas in primates, the most salient cues are received from visual and auditory inputs (reviewed in Behrendt 2011), suggesting distinct fundamental neurobiological substrates. Patterns of activation of oxytocin and vasopressin-like receptors in social contexts, however, suggest conserved neural networks among different species (Johnson and Young 2017). Indeed, vasopressin and oxytocin modulate social recognition in both rodents and primates (Insel 1992, Carter 2014), suggesting common circuitry despite different sensory input modalities. Common neurotransmitter systems, and potentially common downstream circuitry, make the translation from rodent preclinical to the human case possible, despite qualitative differences in the perceptual apparati used by the different genera.

**Table 4.**
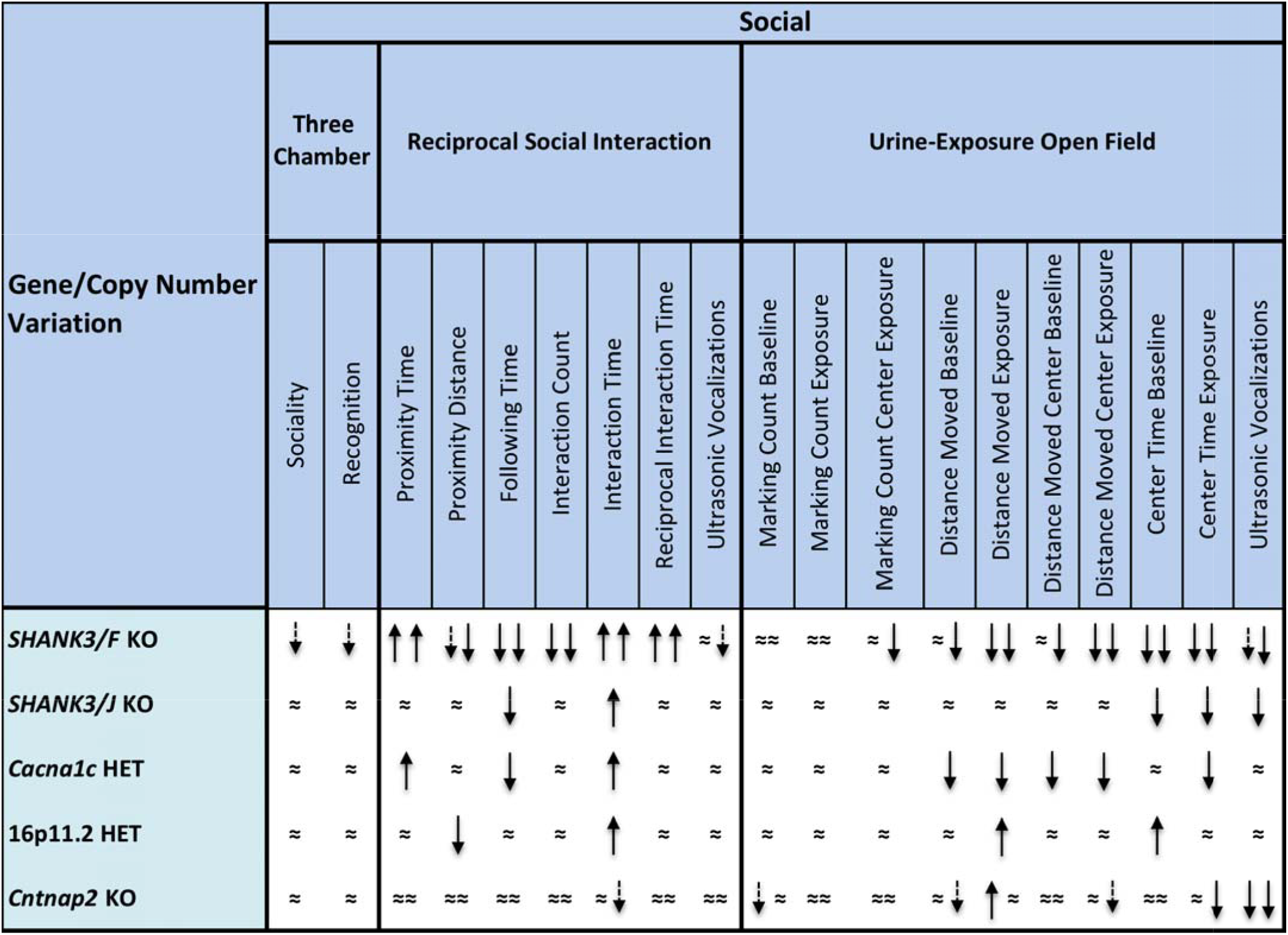
Summary of social behavioral results across the ASD animal models studied. Directions of effects are shown comparing each model system to their corresponding WT controls (upward arrow: increased, downward arrow: decreased, ≈: no difference, dashed shorter arrow: non-significant trend). Two symbols occupying a single cell reflect both the original study and replication results.

Our results point to several difficulties with the reciprocal social interaction test, starting with the fact that time following and interaction frequency differences between mutant and WT mice were always in the opposite direction of interaction time and proximity differences. That is, mice that follow and interacted more also spent less time interacting and remain further away. This pattern is better explained by motor activity rather than by differences in social drive. In human, importantly, social behavior is not a unitary domain and presents differently in different disorders. For example, in Williams and Down syndrome, autism characteristics coexist with hyper-, not hypo-, sociality (Porter, Coltheart et al. 2007, Klein-Tasman, Phillips et al. 2009). Moreover, social behavior patterns are varied and change according to the context (Moss, Howlin et al. 2013). Social behavior seats atop a number of other processes, such as basic and higher-order perceptual processes, emotional states and reactivity, behavioral control, anxiety, etc. Hyper sociality in Williams syndrome, for example, is better explained by decreased behavioral control, and not by either lack of emotion recognition or increased social approach, a finding that required an extensive neuropsychological assessment (Porter, Coltheart et al. 2007). The current approach of modeling all social deficits expected in autism with a single test in mice is clearly an oversimplification and a different approach, one based on efforts to identify underlying constructs and their biological underpinnings, is needed. Uta Frith (2016), argues that a number “startup kits” can and should be recognized in human social behavior: agency recognition (is the agent in front prey, predator, mate, friend, or enemy?), affiliation (recognizing kin, bonding, attachment), alignment (mimicry, resonance, contagion), belonging (identity, trust, loyalty, in group/out group distinction), hierarchy (knowing one’s place, dominance/ submission, alliances), mentalizing (mental state tracking, persuasion, deception, reputation), and morality (fairness, equity, altruism, punishment). Such basic processes, constructs, can be first understood in terms of their underlying biology, and then be back-translated to mice, devising appropriate tests when possible.

##### Juvenile & Adult Ultrasonic Vocalizations

Measurements of USVs in the reciprocal social interaction test showed no significant phenotypic differences across the five model systems, whereas, in the urine-exposure open field test, interesting decreases in the number of USVs emitted were observed in the *Shank3/F, Shank3/J*, and *Cntnap2* KO mice compared to their WT counterparts (see Table 4). These results suggest that USV emission is a measure sensitive to social context in a test not confounded by activity.

#### Conclusions

Despite that many of our findings did not replicate larger effects published in the literature, we found overall excellent replication of most of the results from our previous publications, using the same protocols and animal models, often after consultation with the originating laboratories. Thus, we argue that the problem in the lack of replicability of results resides more in the differences between labs, protocols, husbandry, data handling, and statistical analysis, than in the variability of the animal models themselves.

There has been much recent discussion surrounding the issues of the replicability and reproducibility of preclinical results (ie., Button, Ioannidis et al. 2013, Begley and Ioannidis 2015, Kafkafi, Agassi et al. 2018). We do strongly agree that exploratory results should be confirmed in the same lab through replication but to do this, incentives around funding and publication must be strengthened. Between labs, reproducing confirmed findings with slight environmental and experimental deviations should lead to greater understanding of the robustness of models and associated phenomena. None of this is possible, of course, without complete and transparent reporting of results and, optimally, access to datasets underlying both positive and negative (often unpublished) results along with associated data such as animal health records (i.e., Kilkenny, Browne et al. 2010, Brunner, Balci et al. 2016). The rapid evolution of increasingly sophisticated computational modeling techniques and inexpensive data storage are beginning to converge with the efforts made by several groups to develop a global ontology and common data elements to harmonize massive amounts of preclinical data (e.g., Ferguson, Nielson et al. 2014, Smith, Hicks et al. 2015, Lapinlampi, Melin et al. 2017, Hume, Chow et al. 2018, Wang, Liao et al. 2018). By analyzing data en masse, the hope is that researchers will be able to develop hypotheses on more solid empirical ground (Haefeli, Ferguson et al. 2017) and drug developers can more accurately determine what model systems and tests to pursue; both spending less time and financial resources following independent underpowered studies (see for example Freedman, Cockburn et al. 2015).

In summary, we reiterate our strong belief that mouse model systems of human disease that present with etiological validity (i.e., where there is homology between cause of pathology in both human and animal model system) and construct validity (i.e. there is homology of the pathological process) are fundamental tools for the understanding of gene function, pathophysiology, and are necessary for drug and treatment development. We further argue, based on a complete review of data obtained from three comprehensive studies, that simple measures of known biology such as startle reactivity and self-grooming may provide a venue to link rodent and human pathology. Other measurements, namely social behavior, require refinement and a proven homology to specific constructs underlying the idiosyncratic social profiles described for each syndrome. The face validity approach that supports the use of simple social tests, or anxiety tests for that matter, brings little promise for ASD, an area of research in critical need of robust and replicable preclinical science.

## 6. Acknowledgments

We are grateful to Judy Watson-Johnson, Vanessa Rivera, and Melinda Ruiz for breeding efforts. Thanks to Ian Russell for assistance with all video-based systems and to Mitch Silverstein who offered invaluable support in apparatus development.

