## Supplementary material for "Mouse Model Systems of Autism Spectrum Disorder: Replicability and Informatics Signature": S1 Methods

Replicability in Mouse Models of ASD

**S1 Methods. PCR assays and bioinformatics for SmartCube.**

**Genotyping of the *Shank3/F* Model by PCR assay.**

Genotyping was done by Mouse Genotype (958 Sea Wind Court Carlsbad, CA 92011), based on protocols provided by The Jackson Laboratories.

Tm2G PCR assay:

WT = 482 bp

14080:	TCTAACTCCCAGAGGCCAGA

14084:	AAGGTTGAGCTGGGAGGTCT

KO = ~450 bp

14080:	TCTAACTCCCAGAGGCCAGA

14083:	TCAGGGTTATTGTCTCATGAGC

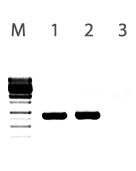

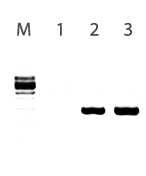

*Figure 1. PCR gel showing a molecular size marker (M) and samples for the Het (1), KO (2) and WT (3) mouse.*

| **Step** | **Temp** | **Time** | **Note** |
| --- | --- | --- | --- |
| 1 | 94^o^C | 3 min |  |
| 2 | 94^o^C | 30 sec |  |
| 3 | 62^o^C | 45 sec |  |
| 4 | 72^o^C | 45 sec | Go to 2, 34 cycles |
| 5 | 72^o^C | 4 min |  |
| 6 | 4^o^C | Hold |  |

*Table 1. PCR conditions for the phenotyping of the SHANK3/F model.*

**Genotyping of the *Cntnap2* Model by PCR assay.**

Genotyping was done by Mouse Genotype (958 Sea Wind Court Carlsbad, CA 92011), based on protocols provided by The Jackson Laboratories.

Cntnap2 WT - 351 bp

13634 (Common):	CTGCCAGCCCAGAACTGG

13635 (WT R):	AGTTGATACCCGAGCGCC

Cntnap2 -/- - ~350 bp

13634 (Common):	CTGCCAGCCCAGAACTGG

10791 (-/- R):	CGCTTCCTCGTGCTTTACGGTAT

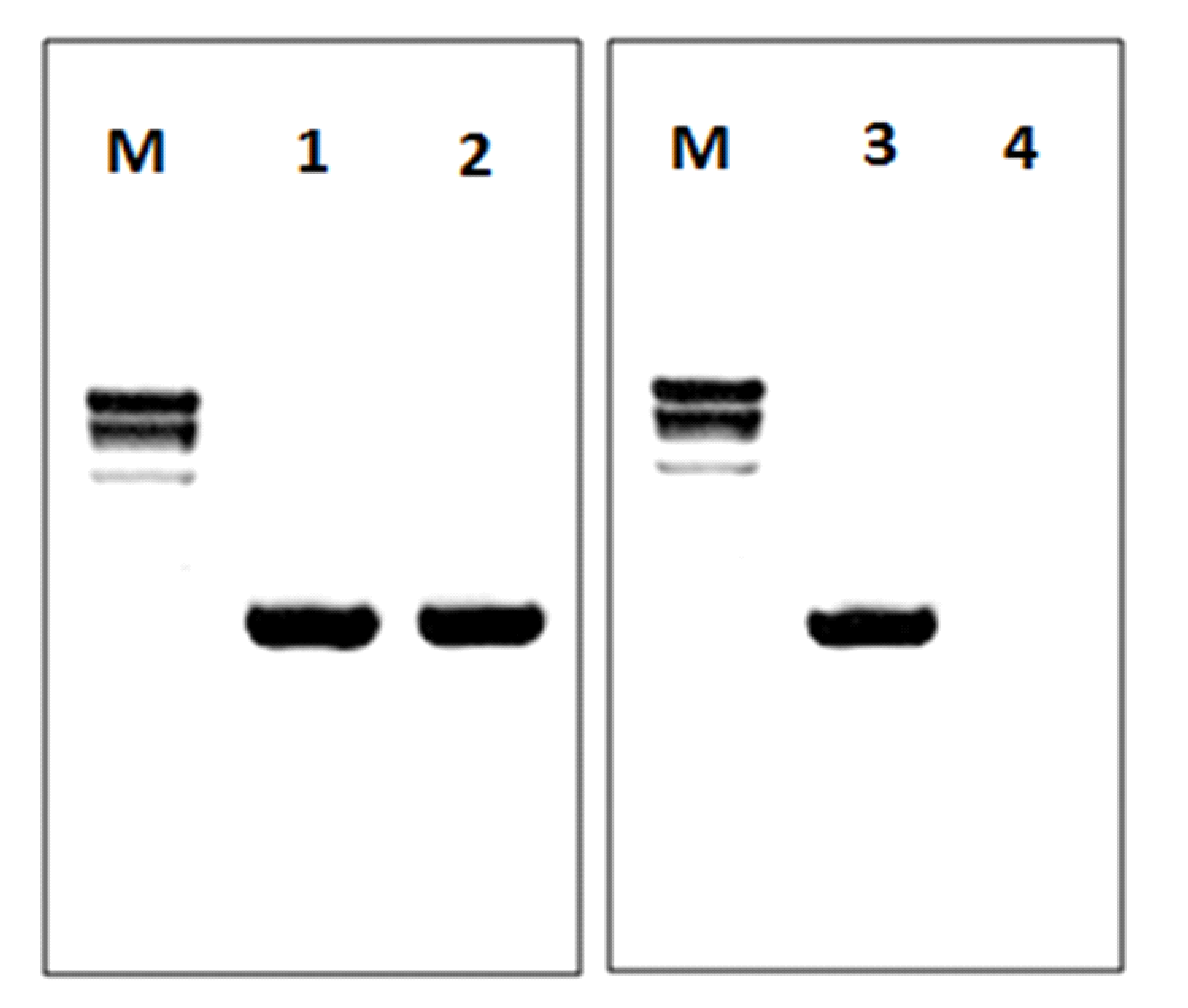

*Figure 2. PCR gel showing a molecular size marker (M) and samples for the WT (1) and -/- (2) mouse.*

| ***Step*** | ***Temp*** | ***Time*** | ***Note*** |
| --- | --- | --- | --- |
| 1 | 94^o^C | 3 min |  |
| 2 | 94^o^C | 30 sec |  |
| 3 | 58^o^C | 30 sec |  |
| 4 | 72^o^C | 40 sec | Go to 2, 35 cycles |
| 5 | 72^o^C | 2 min |  |
| 6 | 4^o^C | Hold |  |

*Table 2. PCR conditions for the phenotyping of the Cntnap2 knockout model.*

**Bioinformatics for** **SmartCube.**

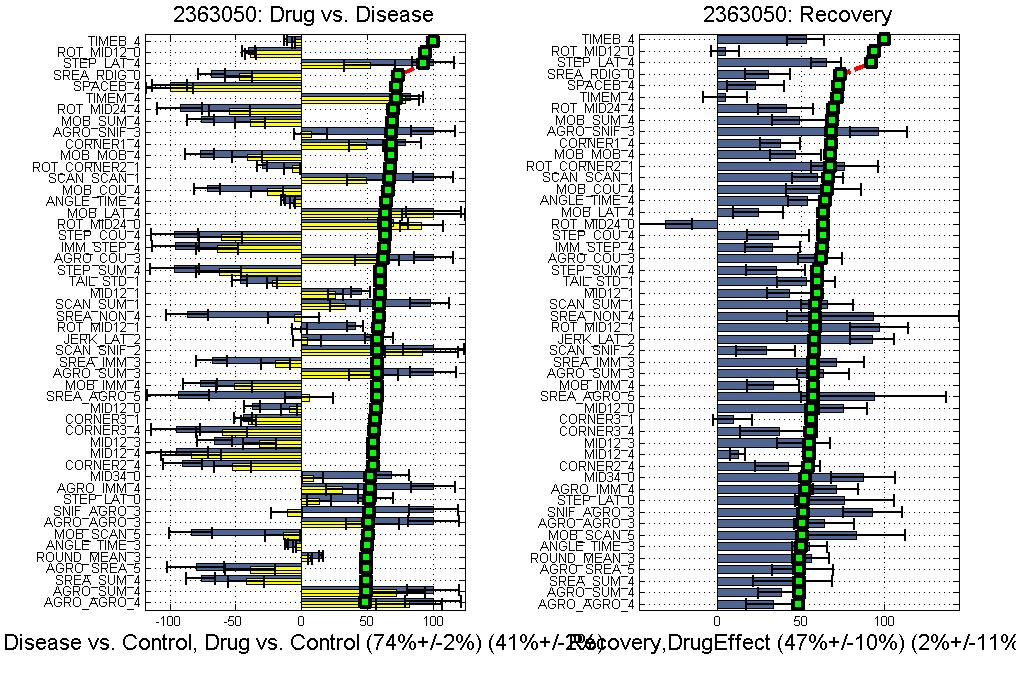

### Feature analysis: de-correlation and ranking. The outcome of any of SmartCube analysis is a vector of hundreds of features (behavioral parameters) that can be used for various analyses (e.g. one run through SmartCube produces more than 1400 features). Many of these features are correlated (e.g. rearing counts and supported rearing counts). Therefore we form statistically independent combinations of the original features (further referred to as de-correlated features) that discriminate between the two groups more effectively. Each de-correlated feature extracts information from the whole cluster of the original features so the new feature space has lower dimensionality.

*Figure 3: SmartCube captures differences in behavior, posture, trajectory, etc. Such features are then processed by the machine learning algorithms which provide a ranked list of features according to which contribute the most to the separation of the two groups. Features can be compared between experimental and control groups, and a % change (increased-shown to the right of zero- or decreased-shown to the left) can then be calculated. In this graph, features that contribute the most are show at the top. Lower ranked features are displayed in order. Relative % feature differences and their ranks (green symbols) can now be displayed on the same graph.*

Next we apply a proprietary feature ranking algorithm to score each feature’s discrimination power (ability to separate the two groups e.g. control and disease). Ranking is an important part of our analyses because it weighs each feature change by its relevance: if there is a significant change in some irrelevant feature measured for a particular phenotype the low rank of this feature will automatically reduce the effect of such change in our analyses so we don't have to resort to the conventional feature selection approach and discard information buried in the less informative features. The ranking algorithm can be applied to either the original or the new features to gain insight about the key control-disease differences (Fig. 3).

### Feature analysis: quantitative assessment of Disease Phenotype. In the new feature space the overlap between the “clouds” (Gaussian distributions approximating the groups of mice in the ranked de-correlated features space) serves as a quantitative measure of separability (or distinguishability) between the two groups. For visualization purposes we plot each cloud with its semi-axes equal to the one standard deviation along the corresponding dimensions (Fig. 4). Note however that while the overlap between any two Gaussian distributions is always non-zero it may not necessarily be seen at the 1-sigma level.

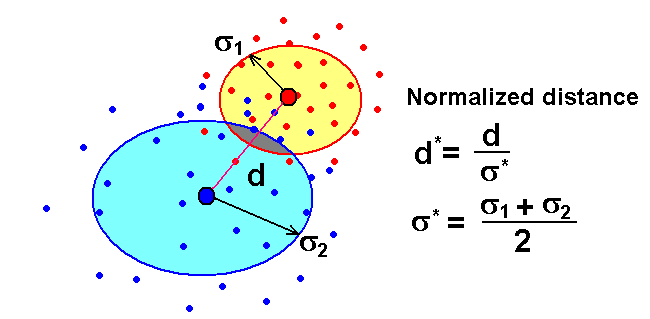
*Figure 4. Visualization of binary discrimination in the ranked de-correlated feature space. The two highest ranked de-correlated features are chosen to form the 2D coordinate plane for visualization purposes. Each dot represents a mouse. Mice from the control group are shown as blue dots and mice from the disease group are plotted in red. The other convenient (from a scale perspective) but equivalent measure derived from the cloud overlap is discrimination probability = 1 - overlap which measures how reliably a classifier can be trained to discriminate between groups A and B above the chance level zero corresponding to 100% overlap and no ability to distinguish the two groups above the chance level whereas 100% meaning the error free discrimination.*

**Significance.** Discrimination significance (generalized p-value) is calculated in the following way. First, each labeled set of candidates is randomly split with 1:3 ratio, where larger groups from each set are used to calculate discrimination probability using methods previously described. This procedure is repeated multiple times with different random splitting for each iteration to build distribution of “true” discrimination probability *p_true_* (step 1). Number of iterations is limited by a fraction of the total number of split combinations available. Next, all candidates from both groups are combined together without individual class labels (step 2). Similarly to the previous step, this set is split randomly multiple times. Larger group of candidates for each split is randomly divided into two “classes” and used to calculate discrimination probability. After many iterations, distribution of “random” discrimination probability *p_random_* is built (step 2). Both distributions are normalized and their mutual weighted overlap is calculated. The resulting value is a generalized quantity of what is well known as p-value of statistical significance.

S1 Table: SmartCube

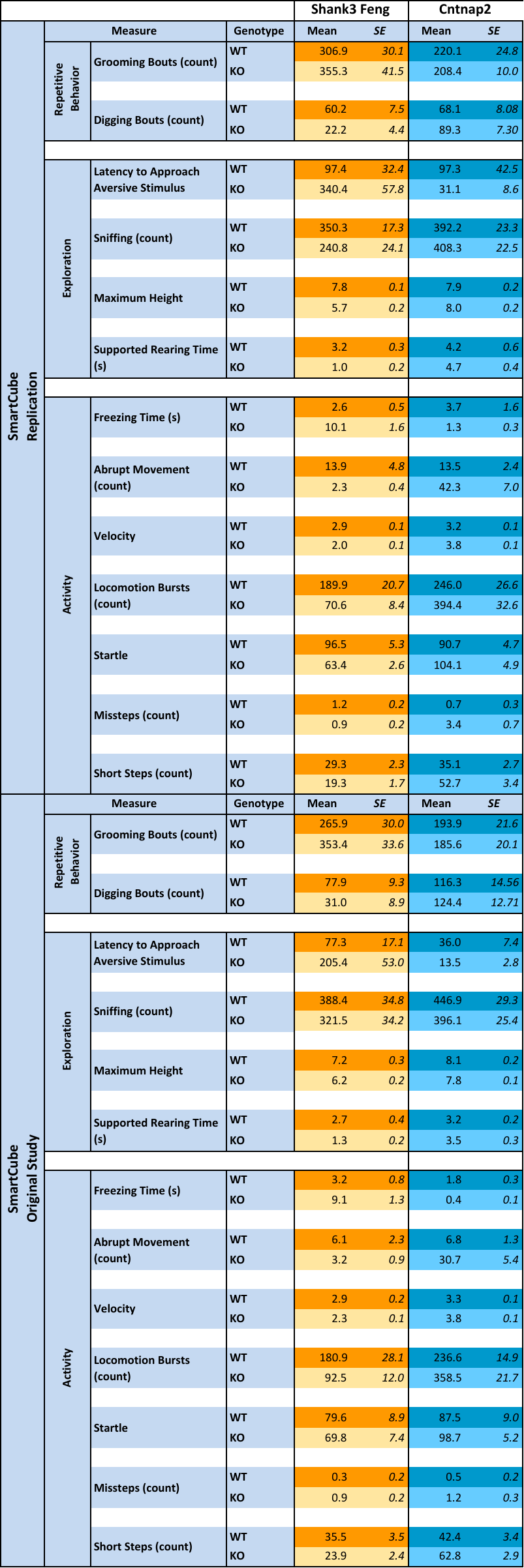

S2 Table: Body weight

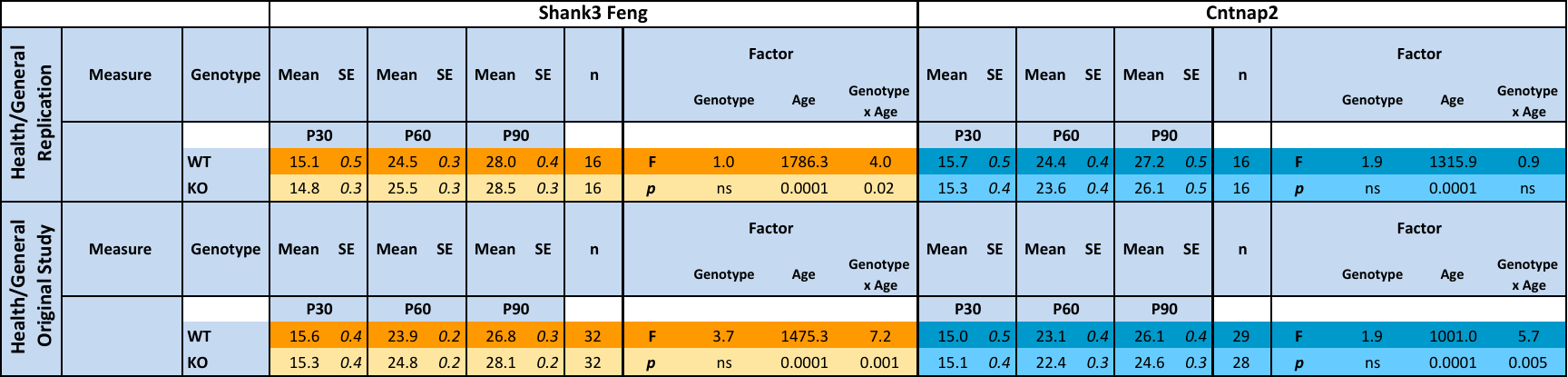

S3 Table: Reciprocal social interaction

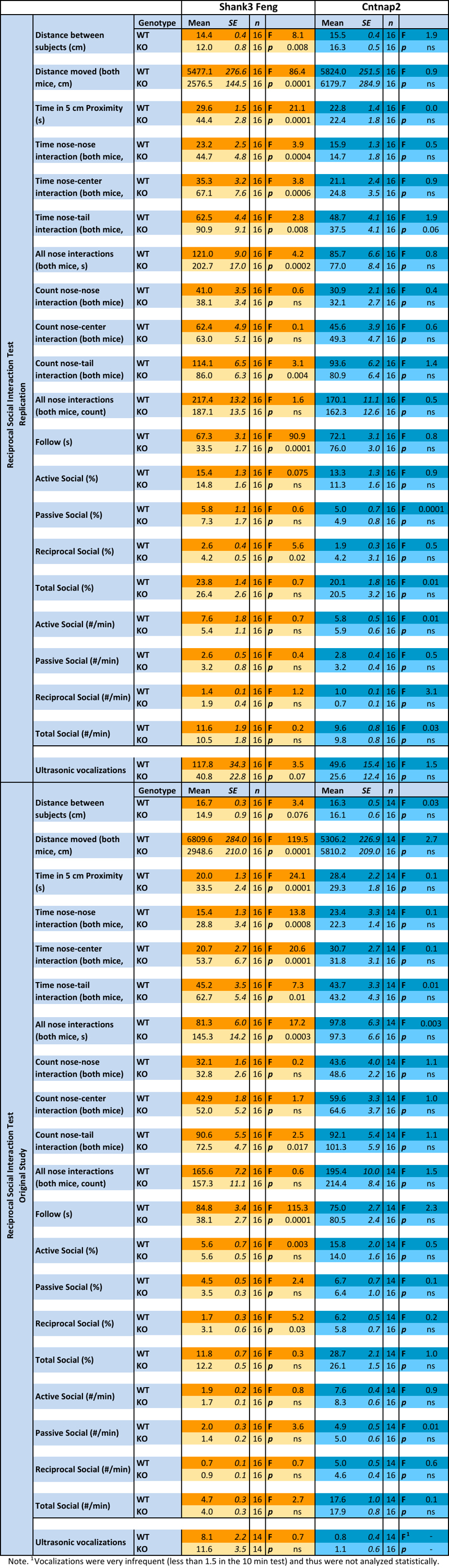

S4 Table: Urine-exposure open field

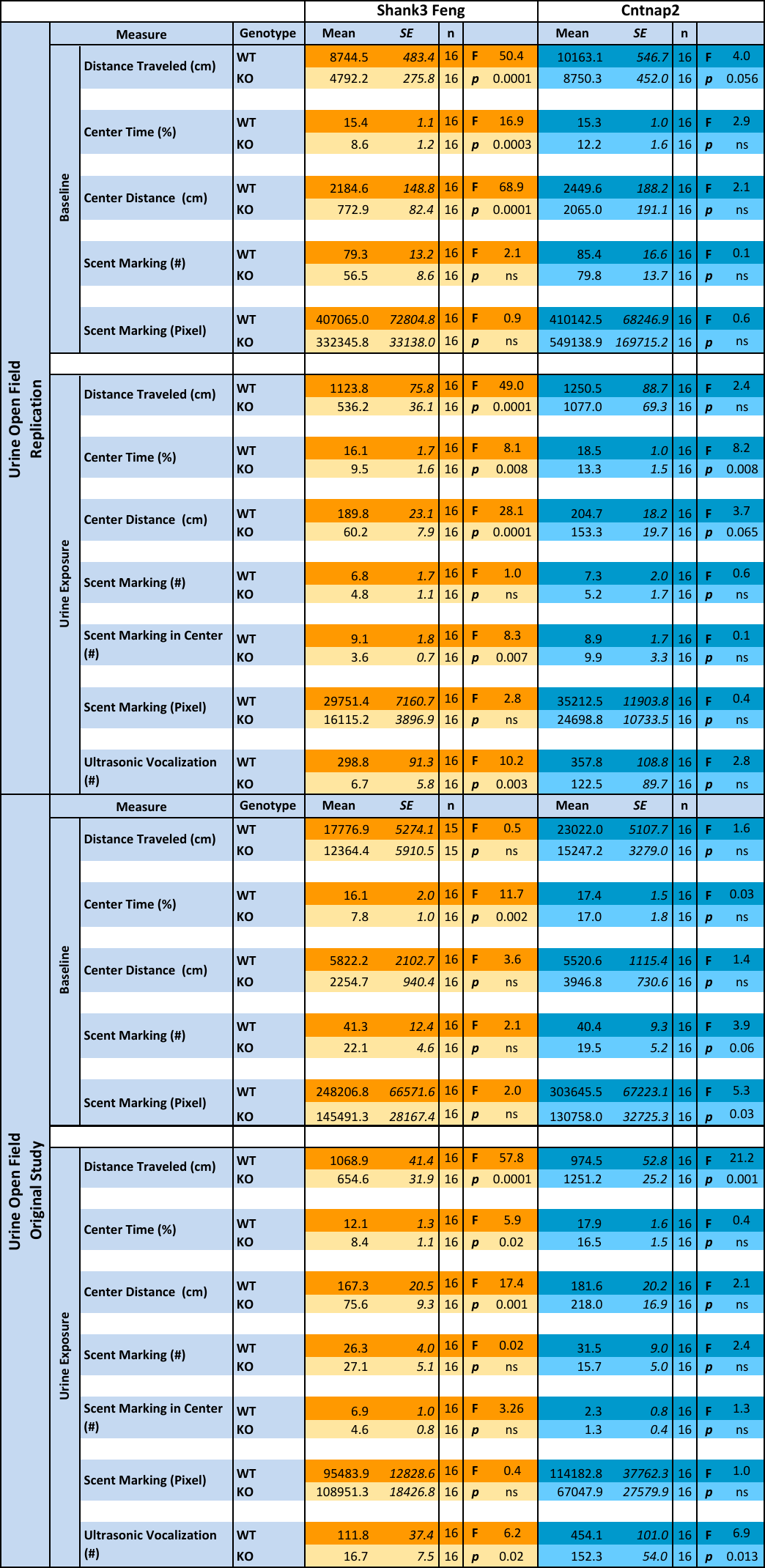

S5 Table: Standard open field

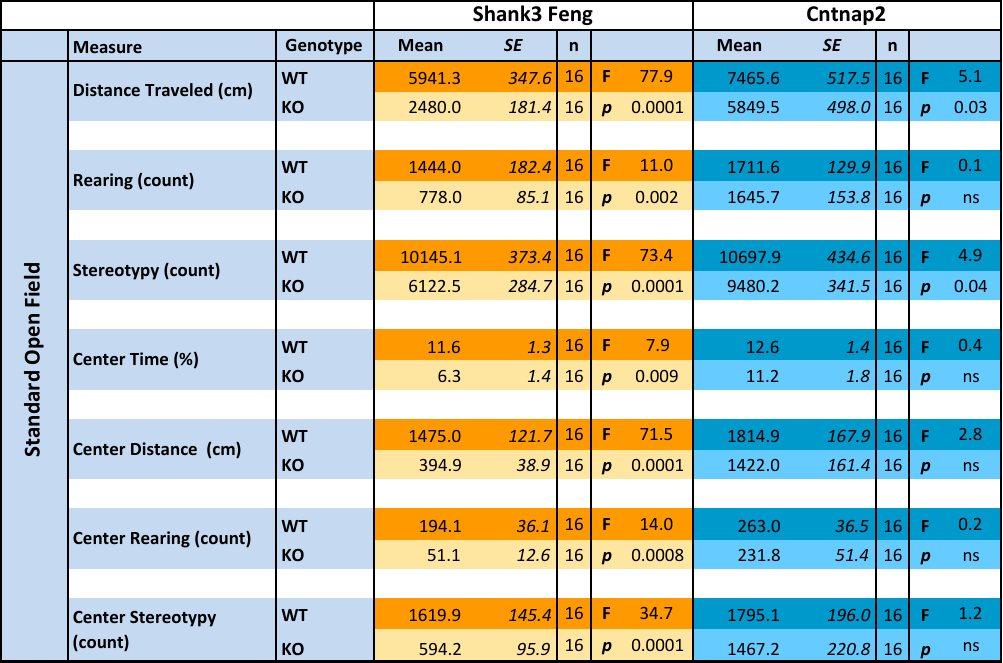

S6 Table: Grooming test

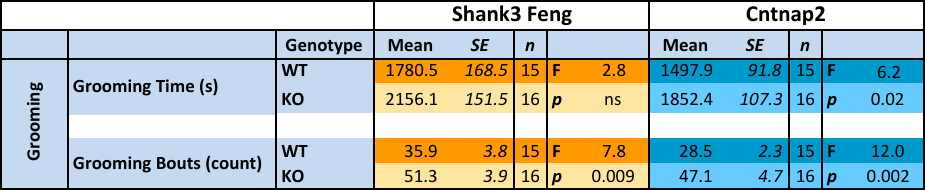
